# Calibration and validation strategy for electromechanical cardiac digital twins

**DOI:** 10.1101/2025.03.06.638897

**Authors:** Zhinuo Jenny Wang, Maxx Holmes, Ruben Doste, Julia Camps, Francesca Margara, Mariano Vazquez, Blanca Rodriguez

**Author notes:** Corresponding author: Zhinuo Jenny Wang Email address, Address: Wolfson Building, Parks Road, Oxford, OX1 3QD, United Kingdom.

## Abstract

State-of-the-art cardiac electromechanical modelling and simulation form the basis for recent developments in cardiac Digital Twin technologies. However, a comprehensive evaluation of electromechanical models at cellular, tissue, and organ level has yet to be performed that addresses both ECG and pressure-volume biomarkers. Such an evaluation would build credibility for applications of cardiac Digital Twins in clinical research and therapy development.

We aimed to follow ASME V&V40 standards for calibration, validation, and sensitivity analysis of ventricular electromechanical models under healthy conditions, to progress towards the Digital Twin vision. We performed a multi-scaled review of ventricular electromechanics and compiled a dataset for calibration and validation incorporating ECG, pressure-volume, displacement, and strain biomarkers.

When applied to a biventricular multiscale model, we achieved healthy calibrated values for the QRS duration (89 ms), QT interval (360 ms), left ventricular ejection fraction (LVEF) (51 %), peak systolic pressure (14 kPa), end diastolic (105 mL) and end systolic volumes (51 mL), peak ejection flow rate (180 mL/ms). Model validation was performed by comparison to displacement and strain biomarkers including systolic atrioventricular plane displacement (1.5 cm), systolic fibre strain (−0.18) and longitudinal strain (−0.15). Sensitivity analysis of model parameters at cellular and ventricular scales was also performed to inform model calibration and as a first step towards uncertainty quantification. We quantified the effects of variability in ionic conductance, mechanical stiffness, cross-bridge cycling dynamics, and systemic circulation on action potential and active tension dynamics at the cellular scale, and on ECG, pressure-volume, displacement, and strain biomarkers at the ventricular scale. Simulations showed that the relationship between healthy LVEF and T wave biomarkers was primarily underpinned by variability in L-type calcium channel conductance and SERCA activity through multi-scale effects. In this study, we provide a systematic framework for credibility assessment of cardiac electromechanical models based on both ECG and mechanical biomarkers, as a foundational step towards the cardiac Digital Twin vision.

## Introduction

Digital Twins are expected to play an increasingly important role in healthcare to enable tailored therapy design for precision medicine. The importance of this emerging technology was attested to by recent regulatory attention and the publication of the ASME V&V40 Standard in 2018^1^. The Standard provided a risk-based framework for evaluating their credibility for biomedical applications. A cardiac digital twin is envisioned as a patient-specific computational model of the heart, personalised from multi-modal clinical data and continuously updated to support diagnosis, prognosis, and treatment planning^2–4^. This is a goal that no published cardiac electromechanical study has yet simultaneously fulfilled, and towards which credible, systematically validated models are an essential first step.

At the core of the cardiac Digital Twin are realistic high-fidelity mechanistic models that can coherently assimilate multi-modal and multi-scale data and can explain disease mechanisms and predict therapy outcomes^5–7^. An important aspect of cardiac function is the coupling of electrophysiology and mechanics in the ventricles^8–10^. To describe electromechanical function, multiscale, mechanistic models are built on mathematical descriptions of electromechanical coupling at cellular, tissue, and organ scales. The biophysical detail gives the model the advantage of high explainability and predictive power over statistical or machine learning approaches. Electromechanical cardiac models have had broad applicability in cardiology, such as in unravelling the relationship between the electrocardiogram (ECG) and mechanical deformation^11^, the ECG and ejection fraction in post-myocardial infarction^12^, demonstrate how mechanical deformation provides triggers and substrate for arrhythmia^13^ and affects re-entrant wave stability^14^, and predicting therapy outcomes in hypertrophic cardiomyopathy^15^ and heart failure^16^. Recent developments towards the Digital Twin vision have enabled increasingly personalised models through assimilation of omics, imaging, blood pressure, and electrocardiogram (ECG) data^9,17–19^ and promises to deliver patient-specific therapy planning^20^ in the near future.

Despite significant developments, a lack of thorough reviews of model credibility considering both electrophysiological and mechanical properties from cell to organ scales hinders wider progress acceptance and impact of cardiac Digital Twins. Thus far, cardiac applications of the ASME V&V40 standards have focused on device optimisation using fluid-structure interaction models for left ventricular assistive devices^21^ and for artificial heart valves^22^, but this has yet to be applied to ventricular electromechanical simulations. Examples of rigorous model evaluation at cellular scale^23–25^ highlight the importance of strictly separating calibration and validation data^23^ and consideration of both electrophysiological and mechanical biomarkers^26^. At the ventricular scale, most studies focused either on the ECG^11^ or pressure-volume biomarkers^27–30^ ^21^ ^31^ but not on both, and deformation and strain biomarkers are rarely considered. Simulation studies have investigated how model parameters relate to specific aspects of cardiac electromechanical function, such as the atrioventricular plane displacement^31^, and how model deformations relate to ECG morphologies^11^, but not in a comprehensive manner. Large scale global sensitivity analyses studies of pressure and volume biomarkers have analysed effects from anatomical variations^28^ and contractility and haemodynamic parameter variations^32^, without any analysis of concurrent ECG biomarker effects. Furthermore, the strategies and data used for calibration and validation in these studies were typically not discussed^8,9,28^. In particular, while Camps et al.^33^ demonstrated personalised ECG calibration in a purely electrophysiological framework, and Strocchi et al.^32^ performed global sensitivity analysis of pressure-volume biomarkers in a whole-heart model, neither study examined the concurrent influence of cellular, mechanical, and haemodynamic parameters on both ECG and pressure-volume characteristics within a single coupled electromechanical framework.

Therefore, in this study we aim to propose a calibration and validation strategy, which starts with gathering the required experimental and clinical dataset for model parameters and simulation outcomes, and considering both ECG and mechanical biomarkers for the evaluation of high-fidelity ventricular electromechanical cardiac models, as the foundation towards the Digital Twin vision. We then apply ASME V&V40 principles to a multiscale, biventricular electromechanical model with clinical image-based anatomy and realistic ECG, with the aim to 1) reproduce healthy pressure-volume, ECG, deformation, and strain dynamics, and 2) investigate multi-scale mechanisms that underpin the relationship between ECG and pressure-volume biomarkers.

## Materials and methods

This study is structured in two parts, following ASME V&V40 principles. First, we establish a general framework for calibration, validation, and sensitivity analysis of ventricular electromechanical models, comprising a systematic review of biomarkers spanning cellular, tissue, and organ scales, and their organisation into non-overlapping calibration and validation sets. Second, we demonstrate this framework through a specific example, applying it to a state-of-the-art biventricular electromechanical model under healthy conditions. This two-part structure is intentional: the framework is designed to be reusable and applicable to any electromechanical model and dataset, while the specific example provides a transparent evaluation of current modelling capability with consideration of both electrophysiological and mechanical biomarkers.

### Model assessment strategy

Following examples of the ASME V&V 40-based analysis applied in the context of cardiovascular science ^21,34^, we devised a calibration and validation strategy for human ventricular electromechanical modelling and simulation frameworks by considering the following:

#### Question of interest

Can the modelling and simulation framework produce healthy physiological electromechanical function at cellular, tissue, and organ levels including both ECG and pressure-volume loop biomarkers?

#### Context of use

Human ventricular electromechanics models provide mechanistic explanations for various disease conditions and evaluate therapeutic interventions. Here we focused on model evaluation under healthy conditions to build the foundations of model credibility for disease and therapy investigations in the future.

#### Quantities of interest

To ensure model relevance to its context of use, the quantities of interest were chosen to be biomarkers with known links to key cardiac diseases, including Torsade de Pointes, myocardial infarction, hypertrophic cardiomyopathy, dilated cardiomyopathy, heart failure, and pulmonary hypertension. Therefore we selected the following biomarkers from the 12-lead ECG (QRS duration, QT dispersion, T wave amplitude, T peak to T end duration, and JT interval), the pressure-volume relationship (left and right end diastolic and end systolic volumes and end systolic pressures, and left ejection fraction, left pressure upstroke velocity, left peak ejection and peak filling flow rate), and from measurements of displacement (end diastolic and end systolic wall thicknesses, atrioventricular plane displacements, systolic percentage volume change, and apical displacement) and strain (systolic global longitudinal strain, radial strain, circumferential strain, and fibre stretch ratio). The specific relevance of each biomarker to cardiac diseases was summarised (Table 2, Results).

**Table 1.** Initial parameter values and literature basis for the choice of variability ranges for the model inputs. Unless stated explicitly, quoted values from the literature have come from previous modelling choices.

| Group | Parameters | Initial value | Literature ranges | Ranges for sensitivity analysis |
| --- | --- | --- | --- | --- |
| Passive mechanics parameters | a, af, as, afs | 0.61 kPa, 1.56 kPa, 0.70 kPa, 0.46 kPa | 0.61, 1.56, 0.70, 0.46 kPa (fitted to human data shear and axial data)<sup>38</sup><br>0.057, 21.503, 6.841, - kPa (fitted to partial shear pig data)<sup>39</sup><br>0.059, 18.472, 2.481, 0.216 kPa (fitted to full shear pig data)<sup>39</sup><br>2.28, 1.685, -, - kPa (fitted to biaxial dog data)<sup>39</sup><br>0.333, 18.535, 2.564, 0.417 kPa<br><sup>40</sup>29/04/2026 08:20:00 AM | [0.057, 0.61]<br>[1.56, 2.15]<br>[0.7, 6.8410]<br>[0.216, 0.46] |
|  | b, bf, bs, bfs | 7.5, 35.31, 33.24, 5.09 | 7.5, 35.31, 33.24, 5.09 (fitted to human data shear and axial data)<sup>38</sup><br>8.094, 15.819, 6.959, - (fitted to partial shear pig data)<sup>39</sup><br>8.023, 16.026, 11.12, 11.436 (fitted to full shear pig data)<sup>39</sup>;<br>9.726, 15.779, -, - (fitted to biaxial dog data)<sup>39</sup><br>9.242, 15.972, 10.446, 11.602<br><sup>40</sup>29/04/2026 08:20:00 AM | - |
|  | Kct | 5000 kPa | 3333<sup>40</sup>, 10 - 1000 <sup>41</sup> | [10, 5000] |
| Active tension parameters | Tref | 120 kPa * 10 | 120 (fitted to human contraction data)<sup>24</sup> | [1200, 2400] |
|  | Cross-fibre contraction | 30% of Tref | 0 – 60%<sup>10</sup>,<br>46% (rabbit heart tissue measurements)<sup>42</sup> | - |
| Pressure volume control parameters | R_LV | 10 kPa ms/mL | 10 – 100 kPa ms/mL<sup>10</sup>,<br>100 kPa ms/mL<sup>40</sup>,<br>114 kPa ms/mL<sup>24</sup> | [10, 110] |
|  | R_RV | 33.3 kPa ms/mL | A third of left ventricular value<sup>24</sup>,<br>2.0 +- 0.8 mmHg min/L (control patient measurements)<sup>43</sup> | - |
|  | C_LV | 1.5 mL/kPa | 1.5 mL/ kPa<sup>40</sup>,<br>20.5 mL/kPa<sup>24</sup> | [1.5, 20] |
|  | C_RV | 4.5 mL/kPa | Threefold of left ventricular value<sup>24</sup> | - |
|  | P ejection LV | 10 kPa | 9.7 -10.4 kPa (human measurements)<sup>44</sup>,<br>9 kPa<sup>24</sup>29/04/2026 08:20:00 AM | [9, 10] |
|  | P ejection RV | 2.3 kPa | 2.3 +- 1 kPa (control patient measurements)<sup>43</sup>,<br>3 kPa<sup>24</sup> | - |
|  | P diastolic cut-off LV | 1.3 kPa | 1.3 [0.93-1.73] kPa (control patient measurements)<sup>45</sup>,<br>0.1 kPa<sup>24</sup> | - |
|  | P diastolic cut-off RV | 1.5 kPa | 1.91 ± 0.8 kPa, pulmonary diastolic pressure<sup>43</sup> | - |
|  | Atrial filling duration LV | 150 ms | ~150 ms at resting cycle length<br>1000 ms (human exercise measurements)<sup>46</sup> | - |
|  | EDP_LV | 1.3 kPa | 1.3 ± 0.1 kPa (control patient measurements)<sup>47</sup>,<br>~1 kPa(control patient measurements)<sup>48</sup>,<br>1 kPa<sup>24</sup>. | - |
|  | EDP_RV | 1.0 kPa | 11.0 ± 6.3 mmHg<sup>43</sup>,<br>A third of left ventricular value<sup>24</sup> | - |
| Pericardial constraint | K_epi | 500 kPa/cm | 20 kPa/cm with 5e-2 kPa*s/cm for viscous term<sup>49</sup>,<br>500 kPa/cm<sup>31</sup>,<br>0 – 100 kPa/cm (20 kPa/cm specifically for baseline)<sup>10</sup> | [20, 500] |
| Microstructure | Transmural fibre angle endo to epi | -60 to 60 in LV, 90 to – 25 in RV, 0 | -60 to 60 in LV, 90 to –25 in RV, 0 to 90 at outflow tract<sup>50</sup> | - |
|  |  | to 90 at outflow tract |  |  |
|  | Transmural sheet angle to radial axis | 0 degrees | -45 endocardium to 45 epicardium (canine measurements)<sup>51</sup>, 45 endocardium to 0 epicardium (canine measurements)<sup>52</sup>, ~45 degrees at diastole, ~0 degrees at systole (swine DTI measurements)<sup>53</sup> | - |
| Heart rate | Cycle length | 1000 ms | 63+-2 beats/min<sup>54</sup>, 72 +- 13 beats/min<sup>55</sup> | - |

**Table 2.** Experimental and clinical datasets for calibration of healthy ventricular electromechanical models, with summary of why each biomarker was important due to their implications in cardiac diseases. Data were presented in the forms: [x, y]95% were 95% confidence intervals, x±ystd were mean and standard deviations, x±zsem were mean and standard error of the mean, [x, y]range were the minimum (x) and maximum (y), x [y, z]IQR were median (x) and first (y) and third (z) quartiles , x (y)IQR were median (x) and interquartile range (y). UKBB indicates values extracted from the UK Biobank. Unit conversions from original source has been done where appropriate to preserve consistency across the entire table. SCD: sudden cardiac death, HCM: hypertrophic cardiomyopathy, DCM: dilated cardiomyopathy, CAD: coronary artery disease, HF: heart failure, HFpEF: HF with preserved ejection fraction, PAH: pulmonary artery hypertension, MI: myocardial infarction, HHD: hypertensive heart disease, LV: left ventricle, RV: right ventricle, EDV: end diastolic volume, ESV: end systolic volume, SVL: stroke volume in left ventricle, SVR: stroke volume in right ventricle, ESP: end systolic pressure, CMR: cardiac magnetic resonance imaging.

| Calibration dataset |  |  |  |  |
| --- | --- | --- | --- | --- |
| Group | Biomarker | Ranges | Data info | Implication for cardiac diseases |
| 12-lead ECG | JT interval (ms) | 313 [294, 335]<sub>IQR</sub><sup>59</sup> | mixed sex, UKBB CMR, n=76,266 | In patients with QRS > 120 ms, risk of Torsade de Pointes<sup>60,61</sup> |
|  | Tpe (ms) | 62 (12)<sub>IQR</sub><sup>62</sup> | mixed sex, UKBB without life threatening arrhythmias, n=51,574 | Risk of SCD<sup>63</sup> |
|  | T wave amplitude (Tpeak) (mV) | 0.15 [0.11, 0.19]<sub>IQR</sub><sup>7</sup> | mixed sex, UKBB normal ventricular mass, n=36,956 | Risk of SCD in HCM<sup>64</sup> |
|  | QT dispersion (ms) | 56.6 [37.0, 84.0]<sub>IQR</sub><sup>7</sup> | mixed sex, UKBB normal ventricular mass, n=36,956 | Severity of CAD<sup>65</sup> |
|  | QRS duration (ms) | 89 [81, 97]<sub>IQR</sub><sup>7</sup> | mixed sex, UKBB normal ventricular mass, n=36,956 | Poor prognosis in DCM<sup>66</sup> and risk stratification for HFpEF<sup>66</sup> |
| Pressure volume | LVEDV (mL) | [88, 161]<sub>95%</sub><sup>35</sup> | female UKBB CMR, n=804 | Predictor of exercise capacity in HFpEF<sup>67</sup> |
|  | LVESV (mL) | [31, 68]<sub>95%</sub><sup>35</sup> | female UKBB CMR, n=804 | Predictor of HFpEF<sup>68</sup> |
|  | RVEDV (mL) | [85, 166]<sub>95%</sub><sup>35</sup> | female UKBB CMR, n=804 | Predictor of functional class in PAH<sup>69</sup> |
|  | RVESV (mL) | [27, 77]<sub>95%</sub><sup>35</sup> | female UKBB CMR, n=804 | Stratifies PAH<sup>70,71</sup> |
|  | SVL (mL) | [49, 100]<sub>95%</sub><sup>35</sup> | female UKBB CMR, n=804 | Incident rate of HF<sup>52</sup> |
|  | LVEsP (kPa) | 16.7 ± 2.2<sub>std</sub><sup>72</sup> | mixed sex, UKBB, n=150 | Associated with hypertrophy<sup>72,73</sup> |
|  | RVEsP (kPa) | 3.2 ± 0.4<sub>std</sub><sup>74</sup> | male, radionuclide ventriculography, n=12 | Stratifies PAH<sup>70,71</sup> |
|  | LVEF (%) | [51, 70]<sub>95%</sub><sup>35</sup> | female UKBB CMR, n=804 | Stratifies HF, MI, and SCD risk<sup>75</sup> |
|  | dP/dt<sub>max</sub> (kPa/s) | 150 ± 30<sub>std</sub><sup>76</sup> | catheter measurement in successful CRT, n=17 | Surrogate measure of ventricular contractility in HF<sup>77</sup> |
|  | Peak ejection rate (PER) (mL/s) | 410 ± 102<sub>std</sub><sup>78</sup> | mixed sex, 3D echocardiography, n=15 | Valvular regurgitation<sup>79</sup> and peak systolic velocity associated with CHD<sup>80</sup> |
|  | Peak early filling rate (PFR) (mL/s) | 361 ± 110<sub>std</sub><sup>78</sup> | mixed sex, 3D echocardiography, n=15 | Points to relaxation dysfunction in CHF<sup>81</sup> |

Cellular biomarkers including action potential duration (APD90), active tension peak (Ta_max_), active tension duration (TaD50) and peak upstroke of active tension (dTa/dt_max_) were not explicitly included in the biomarker list since they have already been used to calibrate and validate the cellular electromechanical model previously^23,26^. However, these properties were evaluated in sensitivity analyses (see below) to inform ventricular simulations.

#### Compilation of biomarker data

The healthy ranges for each quantity of interest (biomarker) were compiled from the literature. Reports from larger datasets were preferred for better estimates of population variability. For this reason, values were preferentially included from the UK Biobank, which is a large-scale biomedical database with health information from half a million UK participants, with more than 40,000 cardiac magnetic resonance recordings available^35–37^. This is then supplemented by smaller studies as needed. The use of gold standard magnetic resonance imaging data was preferentially included over other imaging modalities such as echocardiography for volumetric and deformation biomarkers.

#### Separation of calibration vs validation data

The quantities of interest were separated into a calibration and a validation dataset^23^. The calibration dataset consisted of ECG and pressure-volume biomarkers that clinically defined healthy electromechanical function, while the validation dataset included displacement and strain biomarkers that explored the model’s capabilities for providing mechanistic explanations.

#### Sensitivity analyses towards uncertainty quantification

The model was further validated through sensitivity analyses. To showcase versatility, a wider set of parameters were included in the analyses than has explicitly been linked to any disease or therapeutic target. By varying parameters over biologically informed ranges derived from population variability in the literature, the sensitivity analyses characterise how uncertainty in model inputs propagates to uncertainty in simulated biomarkers, providing a foundation for future formal uncertainty quantification using probabilistic inference approaches.

The following model parameters were included from the cellular electrophysiology (conductances of the fast sodium current GNa, late sodium current GNaL, L-type calcium current GCaL, transient outward potassium current Gto, rapid delayed rectifier potassium current GKr, slow delayed rectifier potassium current GKs, sodium calcium exchanger GNCX, inward rectifier current GK1, sodium-potassium pump current GNaK, peak calcium release Jrel, peak calcium re-uptake SERCA, and magnitude of the stimulus current Istim), the cellular active tension generation (peak active tension T_ref_, calcium sensitivity Cal50, cross bridge transition rates from unbound to pre-powerstroke k_uw_ and from pre- to post-powerstroke k_ws_), passive mechanical properties (bulk stiffness a, fibre stiffness a_f_, sheet stiffness a_s_, fibre-sheet shear stiffness a_fs_, compressibility coefficient Kct, pericardial constraint stiffness K_epi_), and circulatory haemodynamics (aortic resistance R_LV_, aortic compliance C_LV_, aortic ejection threshold pressure P_ejectionLV_).

A review of the literature produced a list of initial model parameter values and their known variabilities (summarised in Table 1). Cellular conductances were uniformly varied from 50 to 200% to cover the experimentally measured ranges as done in previous studies^25^. For passive mechanics, only the parameters carrying units of stiffness were included in the exploration and their initial values were extracted from the non-viscoelastic version of the model fitted to human ex vivo stress-strain measurements^38^ and their ranges extracted from literature. The compressibility parameter (Kct), pericardial constraint stiffness (Kepi), and the active tension coefficient (Tref) parameters were allowed greater ranges of variation since they do not correspond to experimental measurements and their physical meanings were model-dependent. Similarly, the resistance and compliance parameters and the ejection threshold pressure were allowed a large variability range since they were part of a lumped parameter model and do not directly map to experimental measurements.

### A specific example: human biventricular electromechanical model construction, calibration, validation and sensitivity analysis

#### Input data

To demonstrate our strategy, we selected a fully covered biventricular tetrahedral mesh with valvular plugs from the publicly available Rodero et al. database^28^ (female, age 49, weight 90 kg). A fully covered ventricular geometry, extending from apex to atrioventricular plane without truncation, was a prerequisite for applying physiologically realistic basal boundary conditions.

Although four-chamber meshes were available, atrial geometry was excluded to ensure applicability to the most common clinical imaging scenario: cardiac MRI, in which atrial walls cannot typically be faithfully reconstructed. The tools and workflows used to apply the framework to this mesh are openly available at https://github.com/jennyhelyanwe/Cardiac-Electromechanics-Workflow, while the ECG calibration code is based on https://github.com/juliacamps/Cardiac-Digital-Twin, enabling other researchers to apply the same framework to their own patient-specific geometries and datasets.

Since the mesh was constructed based on imaging data at end diastole, the mesh was scaled down isotropically to achieve a diastasis volume of 80 mL^43^ to mimic the process of unloading to diastasis. Fibre *f*_0_, sheet *s*_0_, and sheet-normal *n*_0_ vectors at resting were generated using a rule-based method designed for biventricular geometry with outflow tracts^50^. Ventricular coordinates were taken from the published database^28^, and consisted of the longitudinal coordinate (*uvc_l_*) where *uvc_l_* = 1 at the apex and 0 at the base, a transmural coordinate (*uvc_t_*), where endocardium was *uvc_t_* = 1 and epicardium was *uvc_t_* = 0, and a rotational coordinate (*uvc_r_*), where *uvc_r_* = 0 was in the middle of the left ventricular lateral wall.

#### Human electrophysiological model

Simulation of the electrical excitation and relaxation through the ventricles was through the monodomain model with orthotropic diffusion along the fibre, sheet, and sheet-normal directions. Ionic current dynamics and calcium handling dynamics were modelled using the human ventricular electrophysiological cell ToR_ORd model ^23^, with extensive validation on experimental action potential morphology and calcium dynamics, as well as responses to multi-channel drug block.

Purkinje-myocardial junctions were modelled using a fast endocardial activation layer with isotropic conduction velocity 𝜎_𝑒*n*𝑑𝑜_ on the left and right ventricular endocardial surfaces. Standard 12-lead electrode positions were manually mapped from the heart and torso geometry of a previous study to the current geometry, and the 12-lead ECG was simulated at these electrode positions using the pseudo-ECG method^18^.

#### Mechanical model

The mechanical behaviour was characterised by the balance of linear momentum in a total Lagrangian formulation, considering inertial contributions but ignoring volumetric forces. The reference configuration was the scaled down version of the biventricular mesh (diastasis) 𝛺_0_ ⊂ 𝑅^3^, at 𝑡 = 0. The constitutive law for the passive mechanical behaviour of the ventricular tissue was a nearly incompressible version of Holzapfel and Ogden which was orthotropic in mechanical behaviour^10^. The strain energy density function defining this hyper-elastic continuum is:

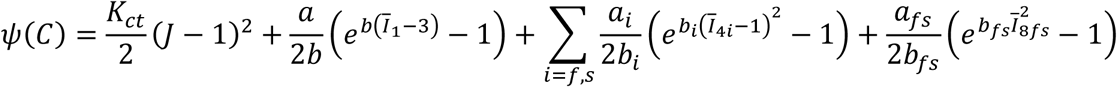

where *J* = *det(F), C* = *F^T^F*, bar notation indicates that 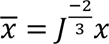, *f*_0_ and *s*_0_ indicate fibre and sheet directions, respectively, in the reference configuration such that 𝐼_1_ = 𝑡𝑟𝐶, 𝐼_4*f*_ = *f*_0_ 𝐶*f*_0_ , 𝐼_4s_= *s*_0_ 𝐶*s*_0_, 𝐼_8*fs*_ = 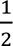 (*f*_0_
𝐶*s*_0_ + *s*_0_ 𝐶*f*_0_), and K_ct_ controls the compressibility of the material. The reader is directed elsewhere^10^ for the precise form of the passive contribution to the second Piola-Kirchhoff stress tensor.

Boundaries were defined as: epicardium 𝛤_𝑒𝑝𝑖_ , left ventricular endocardium 𝛤_𝐿𝑉𝑒*n*𝑑𝑜_, right ventricular endocardium 𝛤_𝑅𝑉𝑒*n*𝑑𝑜_, and the epicardial surface 𝛤_𝑒𝑝𝑖_ ,

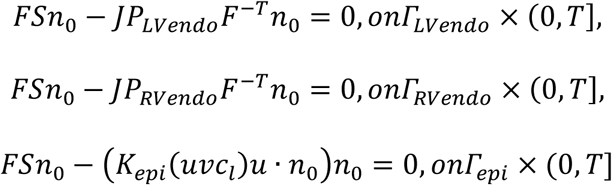

where *n*_0_ was the outward normal defined on the whole material boundary 𝜕𝛺_0_, and 𝐾_𝑒𝑝𝑖_(*uvc_l_*) was the stiffness corresponding to an elastic spring boundary condition applied to the endocardium and was described as a step function of the longitudinal coordinate 𝑙 with magnitude of 𝐾_𝑒𝑝𝑖_ at values of 0 < *uvc_l_* < 0.85 (i.e. applied for 85% of the longitudinal dimension of the heart from the apex). This was a simplified version of the method^31^, which fitted image-derived epicardial displacement to fit a normalised penalty scale that went from 1 from the apex to mid-ventricle, and transitioned through a sigmoidal function to 0 at the base. Rather than imposing the sigmoidal transition, which was highly subject-specific, we imposed a step function instead for the purposes of this study. 𝑃_𝐿𝑉𝑒*n*𝑑𝑜_ and 𝑃_𝑅𝑉𝑒*n*𝑑𝑜_ were the pressures applied to the left and right endocardial surfaces, respectively, and was prescribed by two independent piece-wise functions that describe the five-phases (not including initiation) of the cardiac cycle:

- *Phase 0: Initiation.* Endocardial pressures were linearly increased to reach diastasis values 𝑃_𝑅𝑉𝐷𝑆_ and 𝑃_𝐿𝑉𝐷𝑆_.
- *Phase 1: Active diastolic filling*. Endocardial pressures were linearly increased over time 𝑡_𝐸𝐷𝑃_ to end diastolic values 𝐸𝐷𝑃_𝐿𝑉_ and 𝐸𝐷𝑃_𝑅𝑉_ to mimic the effect of atrial contraction.
- *Phase 2: Isovolumetric contraction*. Electrical activation ensues and active contraction develops. During this, the ventricular pressure was controlled such that the volume was maintained to be constant via:

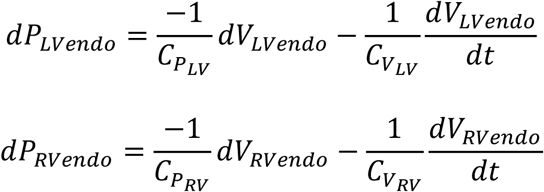

Where 𝐶_𝑝𝐿𝑉_, 𝐶_𝑝𝑅𝑉_ , 𝐶_𝑉𝐿𝑉_, 𝐶_𝑉𝑅𝑉_ , were the inverse of penalty terms for volume difference and volume rates for the left and right ventricles, respectively. The volume rate term was used as stabilisation against spurious oscillations of the ventricular pressure, which might occur due to inertial effects.

- *Phase 3: Ejection.* When the ventricular pressure surpasses the aortic 𝑃_𝐴𝑂_ and pulmonary artery 𝑃_𝑃𝐴_ pressures, the ejection phase begins. To model the blood pressure of the systemic circulation system during the ejection phase, the following two element Windkessel model was used:

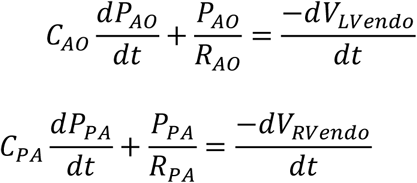

Where 𝐶_𝐴𝑂_ and 𝐶_𝑃𝐴_ were the compliance of the aortic and pulmonary arteries, respectively, and 𝑅_𝐴𝑂_ and 𝑅_𝑃𝐴_ were the resistance of the aortic and pulmonary circuits, respectively. During the ejection phase, the ventricular pressures 𝑃_𝐿𝑉𝑒*n*𝑑𝑜_ and 𝑃_𝑅𝑉𝑒*n*𝑑𝑜_ were modelled as equal to the arterial pressures 𝑃_𝐴𝑂_ and 𝑃_𝑃𝐴_ respectively, disregarding negligible pressure gradients across the arterial valves.

- *Phase 4: Isovolumetric relaxation*. This phase was triggered when the ventricular flow reverses ̇𝑉_𝐿𝑉𝑒*n*𝑑𝑜_ > 0, and ̇𝑉_𝑅𝑉𝑒*n*𝑑𝑜_ > 0, and it follows the same formulation as the isovolumetric contraction Phase 2.
- *Phase 5: Passive filling*. This phase begins when the endocardial pressure drops below the left and right atrial pressures P_LA, P_RA. The pressure was prescribed so that the two ventricular volumes were returned to its diastasis value (DSV_LV, DSV_RV).

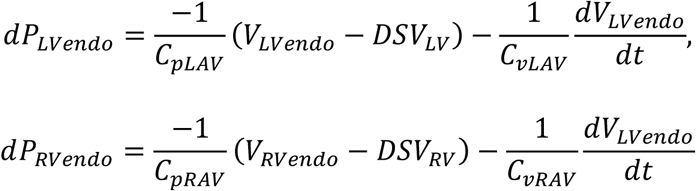

Where 𝐶_𝑝𝐿𝐴𝑉_, 𝐶_𝑝𝑅𝐴𝑉_, 𝐶_𝑣𝐿𝐴𝑉_, 𝐶_𝑣𝑅𝐴𝑉_ were the inverse of penalty terms for volume difference to DSVs and volume rates, for the left and right ventricles, respectively.

The two ventricular volumes 𝑉_𝐿𝑉𝑒*n*𝑑𝑜_ and 𝑉_𝑅𝑉𝑒*n*𝑑𝑜_ were calculated and updated throughout the cardiac cycle by using the divergence theorem and the assumption that the volume was constrained by a flat lid located on the aortic and pulmonary valvular planes, for right and left ventricles respectively, leading to:

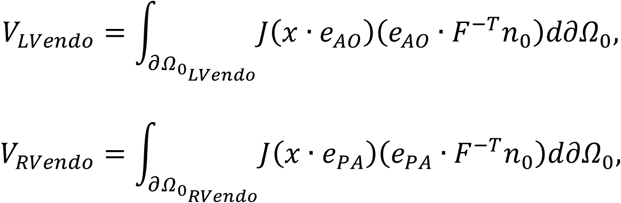

Where 𝑒_𝐴𝑂_ and 𝑒_𝑃𝐴_ were defined so that 𝑒_𝐴𝑂_ · *n*_0_ = 0 on the aortic valve plane, and 𝑒_𝑃𝐴_ ·*n*_0_ = 0 on the pulmonary valve plane.

#### Electromechanical coupling

Electrophysiological activation begins at Phase 2 of the cardiac cycle and drives the active tension generation. This is modelled at the cellular level by coupling of the Land active force generation^24^ with intracellular calcium dynamics from the human cellular electrophysiology ToRORd model^23^, henceforth referred to as the human cellular electromechanical ToRORd-Land model. In order to couple the ToRORd to the Land model, the Hill coefficient of calcium cooperativity and tropomyosin rate constant of the Land model were re-fitted in a previous study^26^ so that physiological active tension could be produced in response to the calcium transient dynamics of the ToRORd model, which were different to the calcium transient to which the original Land model was fitted.

In the Land model, active force generation is modelled using four cross-bridge cycling states (blocked, unbound, pre-power stroke and the force-generating state) and the transition rates between each were calibrated to human skinned myocyte data^24^. The ToRORd-Land coupling is bidirectional, where calcium from the ToRORd binds to troponin C and drives the transition between the blocked and unbound states of the crossbridge system, while the binding of calcium from the Land model acts as a buffer for the ToRORd calcium transient. Furthermore, the length-dependence of troponin calcium affinity/sensitivity in the Land model means that calcium buffering in the ToRORd is dynamically affected by tissue deformation. The Land model was first constructed using skinned myocyte data, then several of the parameters were re-fitted to enable sufficient ejection at the ventricular scale, including the transition rate from the pre-power stroke to force-generating state of the cross bridge (k_ws_), the sensitivity of troponin to calcium binding (Cal50)^24^. At the tissue level, the active tension was then added to the stress tensor to drive mechanical contraction^10^.

#### Literature-informed parameter initialisation

Model parameters were initialised as listed in Table 1. Ionic time constants and conductance values were set to baseline values of the human membrane kinetics ToR_ORd model, which had been calibrated to patch clamp, action potential and calcium transient data^23^. The active tension Land model parameters were as previous calibrated to achieve a physiological calcium transient^26^. The sensitivity analysis ranges in Table 1 were determined following the assessment strategy described above.

#### Verification of numerical solver

Mechanical verification has been treated in more detail in a previous publication^10^. Here, we adopt a set of benchmark problems proposed by Land et al. to verify that the mechanical solver scheme is free from volumetric locking effects. Specifically, we implemented the passive inflation and compared the simulated displacements with the results of simulation across multiple different cardiac mechanics solvers to provide evidence of numerical verification. Details of these benchmarks can be found in the original paper and are briefly explained in Appendix 1. The reader is referred to previous studies for mesh convergence analyses^56^ and sensitivity analyses in an idealized ellipsoid geometry^10^.

#### Calibration to ECG and pressure-volume biomarkers

The 12-lead ECG signals of a female healthy volunteer with similar myocardial mass (108 g) as the biventricular mesh (96 g) was extracted and beat-averaged^33^ for calibration of electrophysiological activation and repolarisation characteristics. The conduction velocities along the fibre and sheet-normal directions were extracted from surgical measurements^57^. The root nodes of earliest activation on the endocardial surface as well as the conduction velocities on the endocardial surface and transmurally were calibrated using the QRS segment of the ECG signal, while spatial heterogeneity in the slow rectifier potassium current (IKs) was calibrated to the ST-T segment of the ECG signal, using a Bayesian-based inference method^18,33^. The calibrated model had a transmural conduction velocity of 48 cm/s, and the fibre and sheet-normal conduction velocities were set to 67 cm/s and 44 cm/s as measured using plunge needle in control subjects^57^, and the Purkinje fibres conduction velocity was set to 300 cm/s^58^. The diffusivity model parameters were tuned to achieve the specified myocardial conduction velocities at the mesh resolution used in simulations.

When the initialised parameter values were used in a preliminary electromechanical simulation, the results undershot healthy values of left ventricular peak systolic pressure by 2 kPa, ejection fraction by 32%, stroke volume by 29 mL, and peak ejection rate by 142 mL/ms, and peak filling rate by 207 mL/ms, and overshot the end systolic volume by 18 mL, and the peak pressure upstroke velocity by 70 kPa/ms. This demonstrated the need for model calibration and explorations of model uncertainties.

Due to the high computational cost of ventricular electromechanics simulations, calibration methods that require high volume of evaluations were infeasible. In this study, we designed a sequential calibration strategy based on knowledge of model sensitivities (described in detail in Appendix 1), targeting multiple quantities of interest in order of clinical important: LVEF, peak systolic pressure, peak ejection rate, and peak filling rate. This involves a series of sampling and parameter selections as follows, where single parameters were uniformly sampled and multiple parameters were sampled using Latin Hypercube Sampling:

1. Sample ejection pressure threshold, Kct, Cal50 and k_ws_ jointly and the parameter set yielding the highest LVEF was selected. These parameters were grouped together because they affect both LVEF and peak systolic pressure simultaneously, through residual active tension, cross-bridge cycling rate, myocardial compressibility, and ejection onset respectively, making independent tuning of each unfeasible.
2. Sample arterial resistance R_LV_ to match peak systolic pressure, since peak systolic pressure is directly dependent on arterial resistance in the Windkessel model.
3. Sample GCaL to further increase LVEF, which consistently undershot the physiology target following Step 2. Even a two-fold increase in GCaL remains within the physiological variability bounds applied in previous studies, and the resulting action potential duration was verified to remain within physiological ranges.
4. Sample k_ws_ again to match peak ejection rate and dP/dt_max_ while maintaining LVEF within target range. k_ws_ is the dominant parameter affecting peak ejection rate and dP/dt_max_ in the sensitivity analysis, and resampling at this stage allows fine-tuning of ejection dynamics without substantially compromising the LVEF achieve in previous steps.
5. Sample diastolic volume change parameter 𝐶_𝑝𝐿𝐴𝑉_ to match peak filling rate.

#### Validation by comparison to deformation and strain biomarkers

The calibrated model was evaluated against displacement and strain biomarkers as listed in Table 2 and not used for model calibration. Atrioventricular plane displacement was calculated by tracking the longitudinal displacement of the top 10% of the geometry using the apex-to-base ventricular coordinate. Wall thickness was calculated using perpendicular projections of the endocardial and epicardial surfaces. Myocardial strains were evaluated at a mid-ventricular short axis slice and at a four-chamber longitudinal, to match DENSE measurements.

Circumferential, radial, and fibre strains were evaluated using the short axis slice, while the longitudinal strain was evaluated using the long axis slice. The strain transients were then time-shifted to begin at the end of diastolic filling (Phase 1) and offset by the end diastolic strain, to match DENSE measurements.

#### Sensitivity analysis at cellular and ventricular scales

A global sensitivity analysis was first performed at the cellular scale using the human cellular electromechanical ToR_ORd-Land model^26^ to identify the key parameters that affect active tension and action potential duration. The conductances of all ionic currents as well as troponin calcium sensitivity and the cross-bridge power stroke rate and myosin head attachment rates (k_ws_ and k_uw_) were varied from 50% to 200%. The key parameters that affect active tension dynamics and action potential duration were identified through ranking the parameters using the total Sobol index. This analysis showed that the key parameters which affected active tension amplitude, duration, and upstroke were GCaL, SERCA, calcium sensitivity Cal50, the cross-bridge cycling rate (k_ws_) and GNaL (Appendix Figure 1).

**Figure 1.**
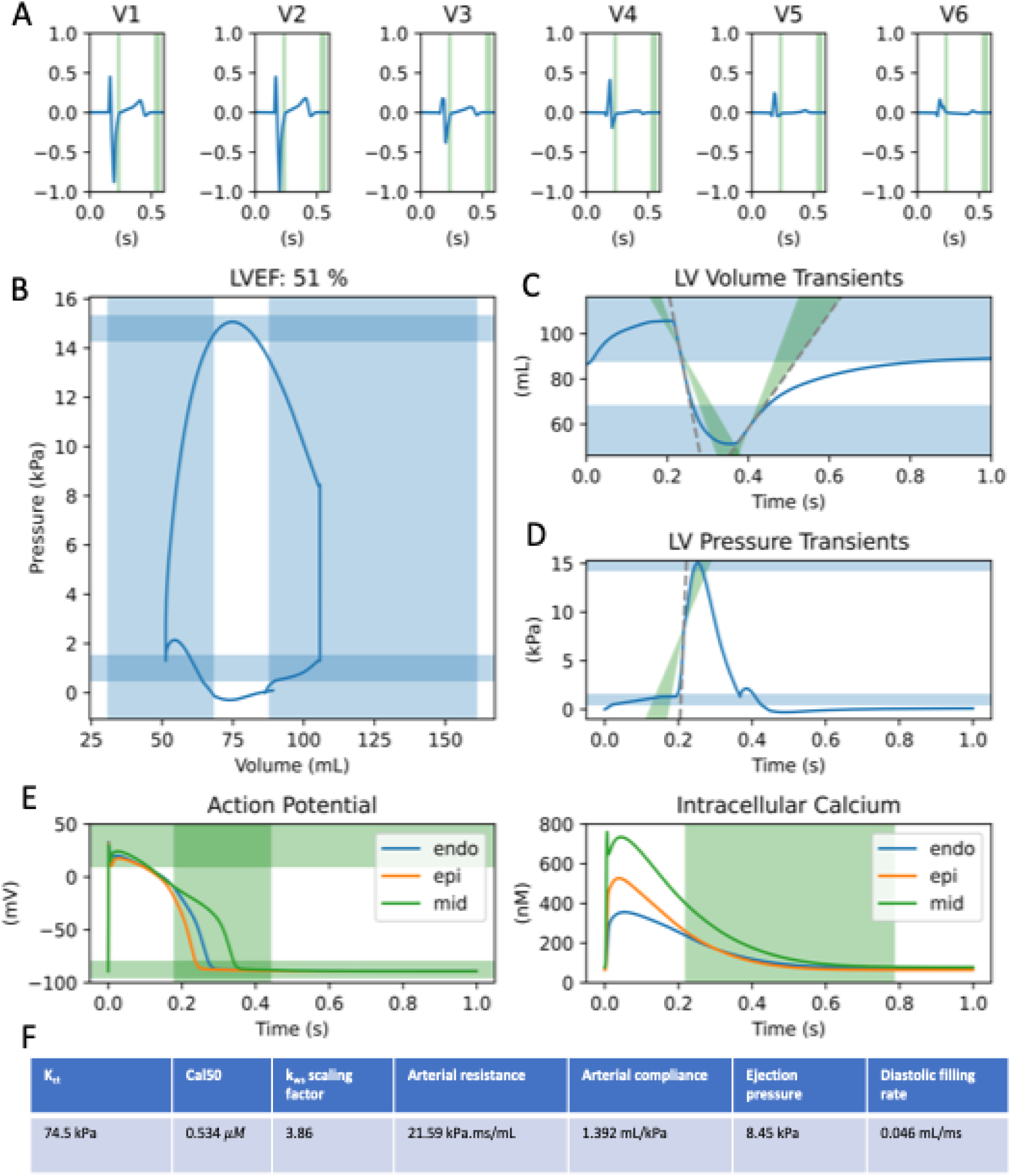
Comparison of (A) simulated ECG, (B) pressure-volume, (C) flow rate, (D) pressure upstroke, and (E) cellular action potential and calcium transient biomarkers of the calibrated baseline electromechanical model with healthy population data ranges shown in green, except for (B, C, D) where the pressure volume ranges were shown in blue. The calibrated model parameters were show in the table (F).

At the ventricular scale, the mechanical and haemodynamic parameters listed in the final column of Table 1 along with the cellular parameters GCaL, SERCA, Cal50, kws, and GNaL were included one-at-a-time sensitivity analyses, varying each parameter uniformly in a total of eight simulations each over the ranges as specified in Table 1 for mechanical and haemodynamic parameters and over 50% and 200% for cellular parameters. Each biomarker as listed in Table 2 were evaluated, and linear regression was performed.

Parameter-biomarker relationships that had p-values less than 0.05 and absolute r-values larger than 0.6 were considered significant and linear relationships. The magnitudes of biomarker effects were normalised by the maximum magnitude for each biomarker. To visualise the relative importance of model parameter on simulated biomarker, a summary diagram was generated with lines linking each parameter to each biomarker. The colour of each line indicated a positive (red) or a negative (blue) relationship. The thickness of each line was scaled by the normalised magnitude of the effect. The transparency of each line was scaled by the r-value as measure of the linearity of relationship (Results Figure 3).

**Figure 2.**
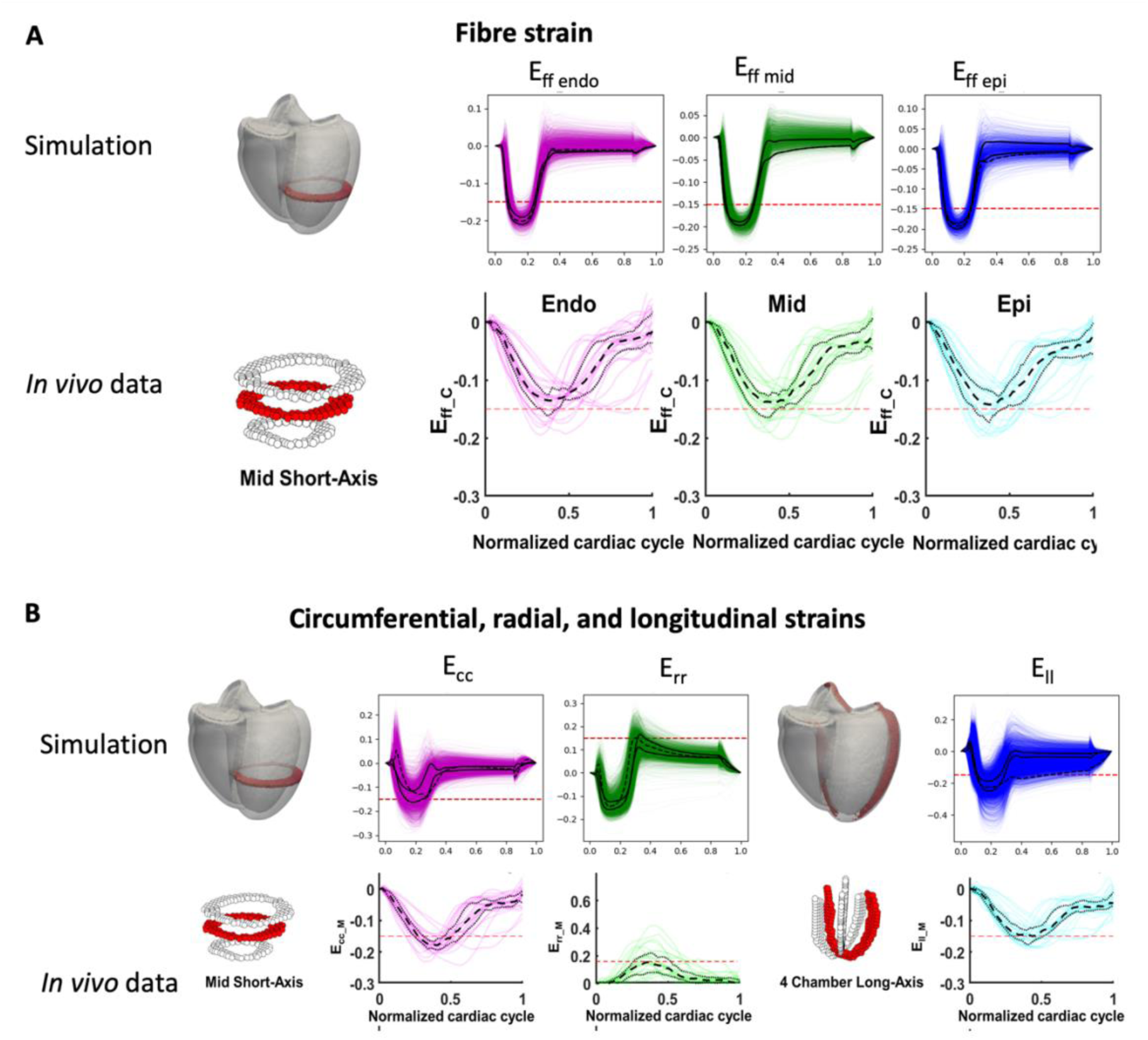
Comparison of simulated baseline left ventricular strains with non-invasive DENSE+cDTI MRI (in vivo) measurements. (A) Comparison of strain along the myofiber direction measured in the mid-short-axis of the ventricles. (B) Comparison of strains in the circumferential and radial directions measured in a mid-short-axis slice and longitudinal strains measured in a four-chamber long axis slice. Median trace is plotted in dotted lines, while upper and lower quartiles in solid lines.

**Figure 3.**
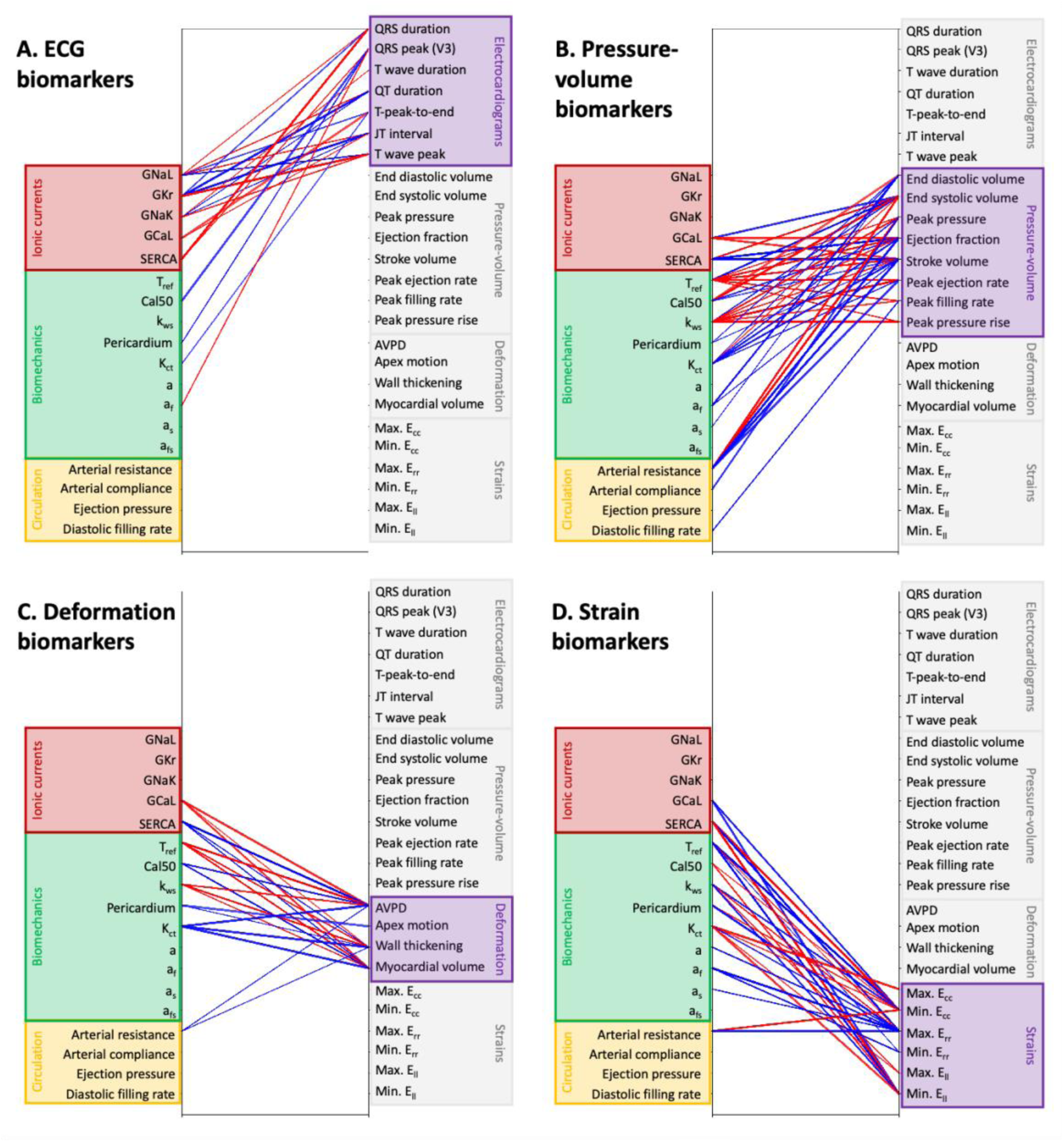
The effect of variability in single model parameters in cellular ionic conductances (red), passive and active biomechanical parameters (green), and circulatory model parameters (yellow) on simulated biomarkers of the ECG (A), pressure-volume (B), deformations (C), and strains (D). Only relationships with p-value <0.05 and r-values higher than 0.6 were plotted. Positive correlations are shown in red, and negative correlations shown in blue. The thickness of each line indicates the normalised magnitude of effect and the width of each line indicates the r-value of the relationship.

### Computations and software

Cellular electrophysiological simulations and Latin Hypercube Sampling were performed using bespoke MATLAB codes. Coupled cellular electromechanics, as well as biventricular electromechanics simulations, were performed using the high-performance numerical software, Alya for complex coupled multi-physics and multi-scale problems^56^ on the JURECA pre-exascale module supercomputer operated by Juelich Supercomputing Centre, Germany, through a PRACE-ICEI (project, as well as on the ARCHER2 supercomputer provided by the UK National Supercomputing Service through the CompBioMedX project (Computational Biomedicine a the Exascale) funded by EPSRC under grant agreement EP/X019446/1. Code for the generation of simulation files for multi-scale sensitivity analysis, calibration and validation, as well as scripts for evaluation of ECG, PV, deformation and strain biomarkers from ventricular simulations can be found at: https://github.com/jennyhelyanwe/Alya_input_setup/

## Results

### Experimental and clinical dataset for model calibration and validation

Table 2 presents the experimental and clinical dataset for calibration of healthy human ventricular electromechanics. For pressure-volume biomarkers and ECG biomarkers, reported values from the UK Biobank^1^ were used, where the number of samples reaches the tens of thousands. Catheter measurements of intraventricular pressure dynamics were rare in the literature and have small dataset sizes.

### Model verification

The simulations of the passive inflation problem set out in the Land (2015) benchmark paper^82^ matched well the observed displacements across different cardiac mechanics solvers (Figure A1). Electrophysiological verification of mesh resolution sufficiency was supported by tuning diffusivity parameters to achieve the specified myocardial conduction velocities, following a previous approach^33^, using an Alya-specific implementation of tuneCV (https://github.com/rsachetto/MonoAlg3D_C/tree/master/scripts/tuneCV).

### Biventricular electromechanics model calibration to ECG and pressure-volume biomarkers

Following model calibration, the simulated ECG showed good R wave progression from V1 to V6 and QRS duration that fell within the ranges shown in Table 1 (Figure 1A) as well as simulated left ventricular ejection fraction of 51% and peak systolic pressure of 14 kPa, with end diastolic and end systolic volumes of 110 mL and 50 mL (Figure 1B, C, D), and a peak filling rate of 310 mL/ms (Figure 1D), all falling within 95% interval of physiological biomarker ranges as listed in Table 1. At the cellular scale, action potential duration, resting membrane potential, peak potential, intracellular calcium transient duration of the calibrated were all within physiological ranges (Figure 1E). The right ventricular end diastolic and end systolic volumes also fell within physiological ranges, though the right ventricular ejection fraction and peak pressure did not, as this was not a key target for calibration. The calibrated model overshot the peak pressure upstroke by 30 kPa/ms and peak ejection rate by 180 mL/ms (Figure 1D). However, the importance of matching to these two biomarkers is lower compared with other haemodynamic markers due to the relative paucity of reported data: n=17 for peak pressure upstroke and n=15 for peak ejection rate, compared with n=804 for ejection fraction and n=150 for peak systolic pressure (Table 2), It is also possible that the calibration could be improved by including some additional model parameters, such as those relating to stretch-rate dependent force generation, even though they did not show a strong effect on single cell active tension upstroke (Appendix Figure A1). These additional parameters might have a stronger effect on the upstroke at ventricular scale than at single cell scale because the single cell simulations were performed under isometric conditions. The calibrated model parameters are listed in Figure 1F.

### Validation by comparison to deformation and strain biomarkers, not used in calibration

Table 3 presents the experimental and clinical dataset for validation of calibrated healthy human ventricular electromechanics. Mechanical displacement and strain biomarkers came from MRI studies with specialised sequences (e.g. DENSE strain imaging).

**Table 3.**
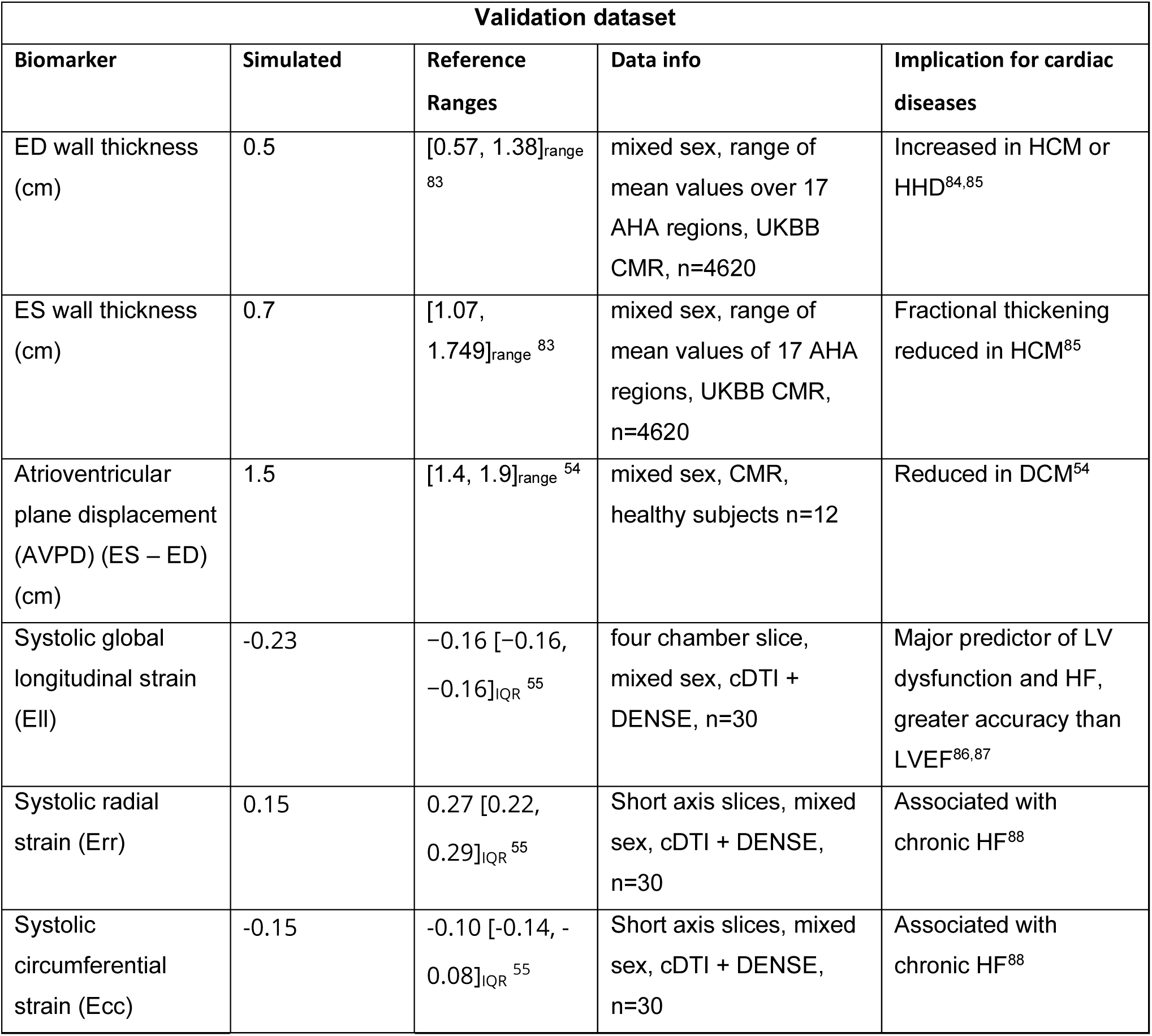
Comparison of simulated validation outputs against reference ranges for deformation and strain biomarkers not used in model calibration. Reference ranges are as reported in Table 2. AVPD: atrioventricular plane displacement, E_ff: fibre strain, E_cc: circumferential strain, E_rr: radial strain, E_ll: longitudinal strain.

Simulation showed broad agreement with the trend and magnitude of *in vivo* strain measurements from DENSE+cDTI images^55^ for fibre strain, circumferential, radial, and longitudinal strain in mid-ventricular and four-chamber view slices (Figure 2, Table 3). Simulated strain traces (n=5004 for short axis, n=9282 for long axis) showed higher variability than *in vivo* traces (n=30). Simulated fibre strain (Eff, Figure 2A) showed agreement in terms of transmural homogeneity when compared with *in vivo* measurements, and the magnitude of fibre shortening was similar to *in vivo* (compare with red line at -0.15 strain). Quantitatively, the simulated average peak strains were E_ff_=-0.20, E_cc=-0.15, E_rr=0.15, and E_ll=-0.23, which fall within or close to the reference ranges reported in Table 2 and showed good agreement in magnitude with *in vivo* measurements (compare with red lines in Figure 2). Radial strain was underestimated relative to the reference range, which is derived from a cohort of 30 subjects and therefore represents a relatively narrow population sample. The timing of simulated peak strains occurred earlier than that *in vivo*. However, this is partially because the model was not calibrated to the cardiac cycle timings of the single individual in the DENSE+cDTI dataset.

In terms of deformations, the simulated systolic vs diastolic atrioventricular plane displacement of the calibrated biventricular (1.5 cm) showed good agreement with ranges reported in the literature (1.4 to 1.9 cm, Table 3). When compared with literature reports of wall thickness changes, simulations showed thickening in systole in agreement with literature. The simulated end diastolic wall thickness (0.5 cm) was lower than the report range [0.57, 1.38] cm (Table 3), and the end systolic wall thickness (0.7 cm) was also lower than the report range of [1.07, 1.75] cm (Table 3). The low wall thickness in both diastole and systole were consistent with the small heart size of the example used (96 g ventricular mass), and a larger and thicker heart would have greater wall thickening and fall better within the population ranges.

### One-at-a-time sensitivity analysis

Figure 3 summarises how variability in single model parameters relate to simulated ECG (Figure 3A), pressure-volume (Figure 3B), displacement (Figure 3C), and strain (Figure 3D) biomarkers. Positive relationships between parameter and biomarker were shown in red and the negative were shown in blue.

ECG biomarkers (Figure 3A) were mostly affected by variabilities in ionic conductances (red block), with some small effects from the mechanical parameters (green block). QRS effects were very minimal, since conduction velocities were not altered in the analysis. QRS duration altered by SERCA (magnitude of change: 2.8 ms), GCaL (1 ms), GKr (1.8 ms), and GNaL (1.3 ms), while normalised QRS amplitude in lead V3 was very weakly affected by calcium sensitivity Cal50 (magnitude of change: 0.04), pericardial stiffness k_epi_ (0.01), and the passive stiffness parameter a_f_ (0.02) (Appendix Figure A1.A). The ST and T wave segments were much more sensitive to model parameters. The QT interval was strongly affected by GKr (magnitude of change: 173 ms), GNaL (92 ms), GNaK (72 ms), GCaL (69 ms), and SERCA (49 ms), with weak effects from calcium sensitivity (Cal50) (1.7 ms). Normalised T wave amplitude was affected by GKr (magnitude of change: 0.1), GNaK (0.06), GNaL (0.06), and SERCA (0.04), GCaL (0.03), with very minor (<0.01) amplitude changes in select precordial leads for Tref (lead V4), Cal50 (lead V4), pericardial stiffness (lead V3) and Kct (lead V2) (Appendix Figure A1.B).

Pressure volume biomarkers (Figure 3B) were mostly affected by the mechanical (green block) and circulatory parameters (yellow block), alongside ionic currents (red block) that affect the calcium signaling system (GCaL and SERCA). Appendix Figure A2 illustrates in detail the parameter effects on the pressure-volume loops. The ejection fraction was predominantly affected by cross-bridge power-stroke rate (k_ws_) (magnitude of change in ejection fraction: 32 %), compressibility (Kct) (16 %), GCaL (15 %), peak active tension (Tref) (12 %), SERCA activity (10 %). There were also some effects on LVEF from GNaL (9%), arterial resistance (6 %), calcium sensitivity (Cal50) (8 %), and very weak effects from pericardial stiffness (4 %), passive mechanical parameters a (3 %), a_f_ (2 %), a_s_ (4 %), and GNaK (2 %), and GKr (2 %). Our models showed that Cal50 not only affected systolic but also diastolic function through altering the diastolic residual active tension, which could contribute to prolonged relaxation in heart failure^89^.

The peak systolic pressure was affected by the same parameters as the ejection fraction, except for GNaK, GNaL and GKr, and with the addition of arterial compliance (magnitude change: 1.1 kPa) and ejection pressure threshold (1.9 kPa). The peak rate of rise of ventricular pressure (dPdtmax) was strongly influenced by only the cross-bridge power-stroke rate (k_ws_) (magnitude of change: 1 kPa/s), with weak effects from calcium sensitivity (Cal50) (0.6 kPa/s) (Appendix Figure A3.A). The peak ejection flow rate (PER) was influenced by Cal50, Tref, k_ws, while the peak filling rate (PFR) was influenced by a larger group of parameters including Tref, Cal50, Kct, GCaL, SERCA, and the bulk parameter controlling the diastolic filling rate (Appendix Figure A3.B).

Mechanical displacement biomarkers (Figure 3C) were predominantly affected by active mechanics (green block) and calcium system conductances (red block), with very minor effects from passive mechanical parameters. Systolic AVPD (Appendix Figure A4.A) was strongly influenced by Cal50 (magnitude of effect: 0.54 cm), GCaL (0.36 cm), Tref (0.33 cm), Kct (0.41 cm), SERCA (0.31 cm), with weaker effect from k_ws_ (0.15 cm), arterial resistance (0.09 cm), and pericardial stiffness (0.08 cm). Systolic wall thickness was predominantly affected by the same parameters as systolic AVPD, except for Kct, which had only a weak effect. Systolic percentage volume change (Appendix Figure A4.C) was strongly affected by the compressibility parameter (Kct) (magnitude of change: 12 %), as was expected, but also strongly affected by GCaL (9 %), Tref (8 %), SERCA (6 %), and kws (3 %). An increase in SERCA and Cal50 shifted the timing of peak volume reduction to earlier during systole.

Simulated strain biomarkers (Figure 3D) were predominantly affected by passive mechanical parameters (green block) and circulatory parameters (yellow block). The influence of parameters on the radial, circumferential and longitudinal strain patterns (Appendix Figure A5) followed broadly that of the AVPD and wall thickness. Of note was that at high values of the ejection pressure, the mid-ventricular circumference expands rather than contracts during systole and fails to return to the reference condition even after relaxation, pointing to failure of proper pumping function (Appendix Figure A5.A).

### Multi-scale mechanisms underpinning ECG and LVEF relationship

Simulation results indicated that the link between ECG biomarkers and LVEF was chiefly underpinned by GCaL and SERCA, which had strong effects on both T wave biomarkers and the pressure-volume loop (Figure 3). Mechanical parameters T_ref_, calcium sensitivity (Cal50), compressibility (Kct), and fibre stiffness (a_f_) had much stronger effects on LVEF than on ECG signatures. On the other hand, ionic conductance parameters GNaL, GKr, GNaK had much stronger effects on the T wave than on LVEF.

Figure 4 illustrates the multi-scale mechanisms of GCaL’s effect on both LVEF and ECG. A four-fold increase in GCaL caused an eight-fold increase in cellular active tension peak and a 200 ms prolongation of active tension duration (Figure 4A), which resulted in a 15 mL decrease in end systolic volume and 2.5 kPa higher peak systolic pressure at the ventricular scale (Figure 4C). GCaL’s dramatic effect on end systolic volume was facilitated by 10 % higher systolic myocardial volume compression, 4.5 % higher systolic longitudinal contraction (Figure 4E, dotted lines), 3.5 % higher systolic circumferential contraction, and 0.4 cm lower (towards apex) systolic position of the atrioventricular plane (Figure 4B). The increase in longitudinal contraction was achieved by a greater downward motion of the basal plane while the apex stayed stationary due to the pericardial constraint. The increase in GCaL caused an increase of 60 ms in cellular action potential duration (Figure 4A), which prolonged the global repolarisation time (Figure 4D) and led to the prolongation of the QT interval (Figure 4C). GCaL did not have a significant effect on the activation pattern and conduction velocity in our simulations (Figure 4D) or on the QRS complex of the ECG (not shown).

**Figure 4.**
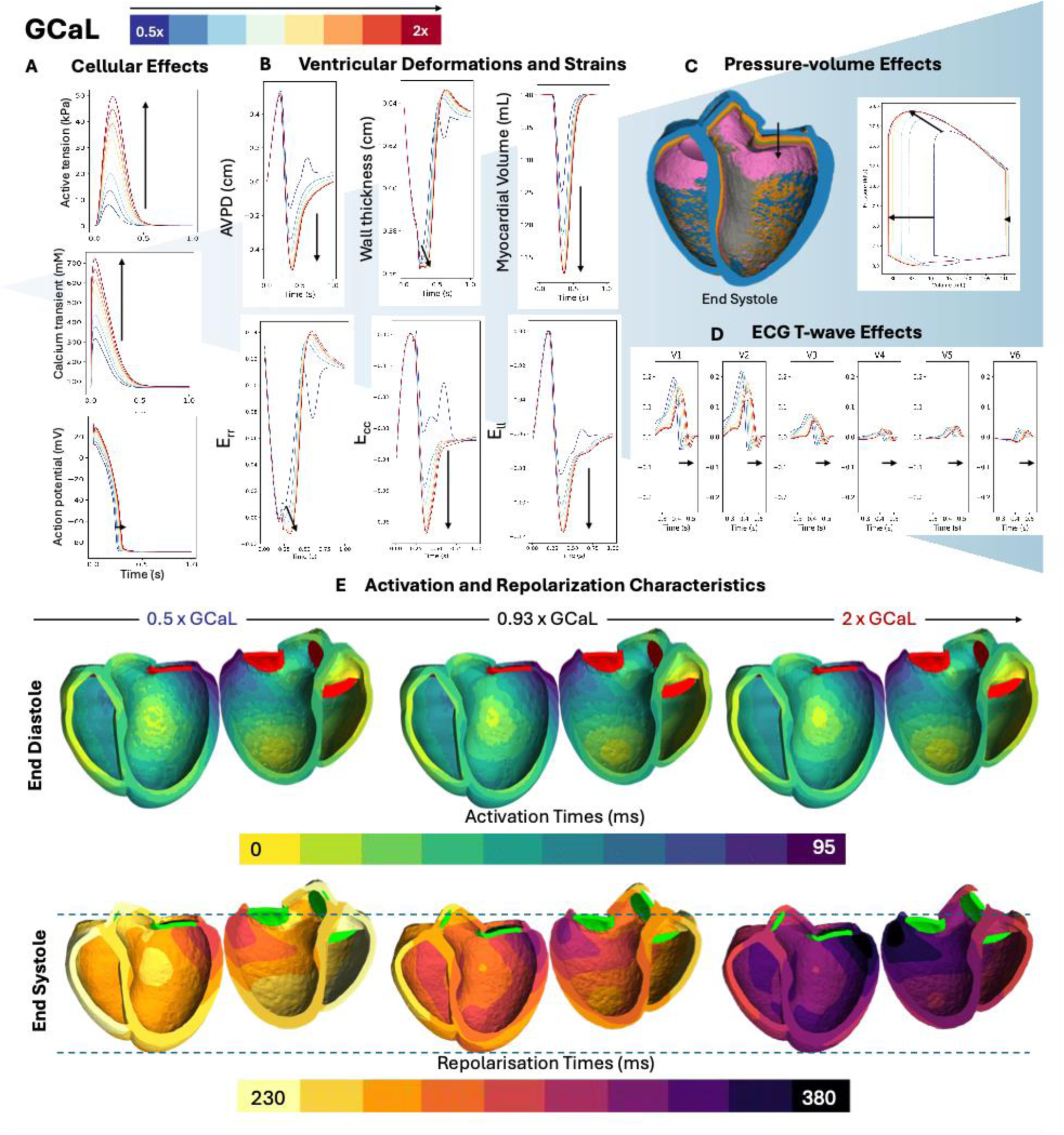
Multi-scale effect of GCaL on active tension, action potential duration (A), ventricular deformation and strains (B), the pressure-volume loop (C) and precordial ECG leads (D). ECG characteristics were explained by the activation and repolarization maps (E), with dotted lines highlighting increased longitudinal shortening with increasing GCaL by the basal plane moving towards the apex while the apical position remains unchanged, due to the presence of the pericardial constraint. Scaling factor of 0.93 was the closest scaling factor to 1 when uniformly sampling eight datapoints in range [0.5, 2].

Figure 5 illustrates how a four-fold increase in SERCA activity caused a 12% decrease in LVEF, with non-monotonic effects on the QT interval. The increase in SERCA conductance increased diastolic residual active tension (Figure 5A), which was sufficient to cause 3 mL reduction in end diastolic volume (Figure 5C). Even though increasing SERCA increased peak active tension at the cellular scale by 9 kPa (Figure 5A), it reduced peak systolic pressure and LVEF at the ventricular scale (Figure 5C). This was because increasing SERCA hastened the arrival of peak active tension by 50 ms and reduced its duration by 100 ms (Figure 5A), which meant that the mechanical contractions were not given enough time to developed, resulting in earlier and weaker systolic contractions (Figure 5B). On the electrophysiological side, while SERCA did not elicit strong APD changes at the single cell simulation, when combined with non-isometric conditions in ventricular simulations, it caused action potential prolongation at values higher than 1.5-fold and resulted in prolonged repolarization (Figure 5E).

**Figure 5.**
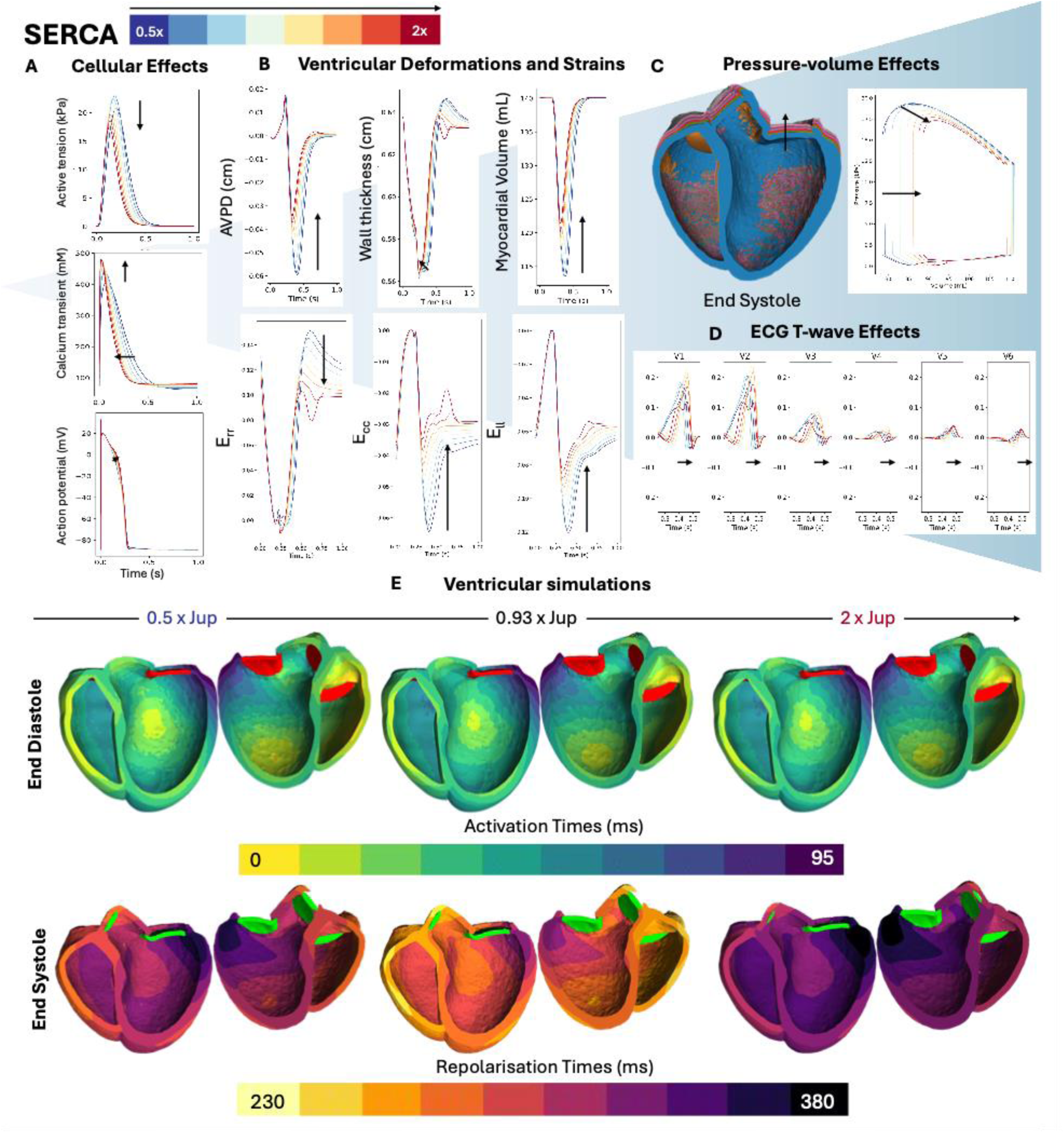
Multi-scale effect of SERCA conductance on active tension, calcium transient, action potential duration (A), ventricular deformation and strains (B), the pressure-volume loop (C) and precordial ECG leads (D). ECG characteristics were explained by the activation and repolarization maps (E). Scaling factor of 0.93 was the closest scaling factor to 1 when uniformly sampling eight datapoints in range [0.5, 2].

## Discussion

In this study we presented the calibration, validation and sensitivity analysis of human healthy ventricular electromechanical modelling and simulation, following the ASME V&V40-based assessment guidelines. The first step was compiling parameters and biomarkers values from multi-modal data characterizing pressure-volume, ECG, displacement, and strain behavior of human healthy ventricles, and separated the data for calibration and validation as well as identify model parameter ranges for sensitivity analysis. The calibration was informed by unravelling the interlinking relationships between simulated biomarkers and model parameters, especially highlighting the plethora of model parameters that affect the ejection fraction. We demonstrated multi-scale mechanistic explanations of the relationship between LVEF and ECG underpinned by variability in L-type calcium channel conductance and SERCA activity. Through considering biomarkers at multiple scales and across both electrophysiology and mechanics, we aimed to provide a transparent audit of current capability by identifying where confidence is established and where further development is needed, as a firm foundation for future refinement Previous examples of applying the ASME V&V40 framework for cardiac model credibility assessment have focused primarily on evaluating computational models of medical devices such as the left ventricular assist device^21^ and artificial heart valves^22^, where model validation was performed by comparing to experimental recordings specifically designed for this purpose. These studies, as well as other examples of model validation without explicit reference to the ASME Standard, commonly have narrow focuses on specific diseases and/or therapies, and the biomarkers used for model evaluation were similarly limited in scope. In the case of disease models, calibrations were commonly performed at the cellular or tissue scale and validation was performed at the ventricular scale through comparisons with known ECG or ejection fraction phenotypes^12,15^. While these studies have been valuable for providing model credibility in their specific disease/therapeutic contexts of use, they have not been designed to give credibility to the underlying computational electromechanics framework. In this study, we instead focused on building a comprehensive calibration and validation strategy for baseline healthy biventricular electromechanics.

We showed in this study that literature values for model parameters do not automatically provide good baseline simulations. We proposed a sequential calibration strategy based on knowledge of model sensitivities, as detailed in the methods. This approach can be adapted for more sophisticated numerical techniques involving the use of emulators and Bayesian statistics to enable faster and more comprehensive explorations of the parameter space^27,29,90^, and enable matching to more stringent calibration criteria such as the inclusion of cellular biomarkers to ensure simultaneous multi-scale compliance. Non-uniform sampling of the parameters based on knowledge of parameter-biomarker relationships could also improve its efficiency.

Our sensitivity analysis results showed that in the simulated ECGs, the T wave was far more sensitive to electrophysiological parameters than to mechanical ones, within the ranges of variability expected from literature reports of each parameter. In our simulations, the ionic conductances GNaK and GNaL primarily affected the T wave through altering the action potential duration (Appendix Figure A1) while having minimal effect on mechanical deformation or strain. GCaL and SERCA strongly affected the repolarization pattern and deformation and strain patterns during systole (Figures 4 and 5). However, other parameters that had a strong effect on deformation and strain, such as Tref, Cal50, pericardial stiffness and Kct had only a minor effect on the T wave (Appendix Figure A2). Previous studies on this question focused on comparing either end diastolic versus end systolic geometry^11^ or a static versus dynamic geometry^91^ when simulating the T wave. These simulations involved a much larger perturbation in deformation than relevant to physiological situations and showed a larger effect on the T wave amplitude than our simulations. Within the context of physiological systolic deformations, however, our results showed that deformation uncertainties had only minor effects on T wave amplitude.

Our simulations showed only minor changes to QRS amplitudes in response to variation in those parameters that influenced the end diastolic geometry: calcium sensitivity, through altering diastolic residual active tension, pericardial stiffness, through restricting diastolic inflation, and the passive mechanical parameters a and af, through altering bulk stiffness and stiffness along the myofibre direction. However, it is possible that the magnitude of changes to the QRS was underestimated in our simulations, because the local activation times on the endocardial surface in our simulations were fixed. This meant that our simulations did not explore the effect of variations in end diastolic geometry on the conduction velocity of the fast endocardial activation layer or on Purkinje conductivity. It should also be noted that other parameters that affect the activation pattern, such as GNa and conduction velocities, were not included in this analysis.

The fact that the initialised literature values of model parameters failed to achieve LVEF above 50 % pointed to potential limitations in our modelling approach. Our final calibrated model required a ten-fold scaling factor on the peak active tension transient, as was found to be necessary in a previous sensitivity analysis study^10^. This is a significant deviation from *ex vivo* measurements of myocardial force production, which is better understood through comparison with similar studies in the field. Strocchi et al. (2023)^32^ used a four-chamber model and required a considerably more modest adjustment of active tension, likely reflecting differences in model geometry, pericardial constraint, and circulatory model.

Gerach et al. (2021) ^9^ achieved stroke volumes close to MRI data but report that systolic pressures in. both ventricles were too high for a healthy heart, attributing this to the steep increase in stress due to their tension model, and similarly report elevated peak ejection rates compared to MRI measurements. They also report that atrial contraction contributes a significant fraction of end-diastolic volume, a contribution absent from biventricular-only models such as ours and a primary likely factor in the compensating active tension required. Zingaro et al. (2024)^92^ found that no single contractility parameter configuration simultaneously achieved physiological peak ejection rate and LVEF, consistent with our own experience. Together these comparisons suggest that the absence of atrial mechanics is a key contributor to the elevated active tension scaling in our model.

Furthermore, this could be explained by other model limitations, including volumetric locking effects, even though benchmarking our mechanical solver as in Land (2015) ^1^ did not show volumetric locking effects. Further analysis is, however, warranted against a more comparable elastodynamic systems with orthotropic passive mechanics, such as presented in the recent Arostica (2025) benchmark problems^93^. Other model simplifications could also contribute to the difficulty in achieving a healthy LVEF. In our simulations, we saw that the LVEF was strongly sensitive to changes in the incompressibility of the tissue (Kct), such that an increase in compressibility of the myocardial tissue helped to increase LVEF. The systolic volume has been reported to change up to 13%^1^, pointing to limitations of assumptions of incompressibility in the modelling of the myocardial tissue and ignoring poro-elastic^95^ and visco-elastic effects^38^. This is further highlighted by the fact that our model did not achieve a high amount of wall thickening in validation, which meant that the high compressibility required in our model to achieve LVEF could be compensating for a lack of sufficient wall thickening, in addition to anatomical effects. Previous experimental and simulations studies have shown how wall thickening is related to sliding of sheetlets^96,97^. However, in our simulations we did not see a significant effect of the sheer stiffness parameter a_fs_ on wall thickness or LVEF. Another possible avenue of exploration would be the effect of sheet and fibre orientations on the thickening effect, as explored in a previous simulation study^97^.

Displacement and strain biomarkers (Figure 2D) were far more sensitive to calcium handling dynamics, cross-bridge cycling rate, and contractility than to passive stiffness parameters, which indicates they would be better indicators of systolic rather than diastolic dysfunctions in HF. The sensitivity of strains and displacements to the pericardial stiffness parameter was unsurprising due to the association of reduced strains in diseases affecting the pericardial sack^98^. The sensitivity of longitudinal and radial strain to the incompressibility of the myocardium (Kct) mirrors the sensitivity of the LVEF to that parameter, which was in accordance with the fact that those two strains were surrogate markers of LVEF and show better predictive power^86,87^.

### Limitations

While the electromechanical model presented is advanced in terms of its human-relevance and multi-scale complexity, it contains modelling simplifications that can be extended in future work to allow its application to specific areas of investigation.

The model uses an isotropic downscaling of the end diastolic biventricular anatomy to approximate the unloaded reference geometry, to achieve a physiological diastasis volume. While our simplification allowed us to match population-based volume metrics, more physically accurate methods of unloading the geometry^1,2^ will be necessary if the aim is to personalise mechanical stiffness parameters^100–102^ and investigate diastolic dysfunction^103,104^. Moreover, even in the unloaded state, the myocardial tissue is not without residual strains^105,106^ , which can be important to take into account when inferring patient-specific diastolic function.

Our lumped parameter model of the circulatory system was sufficient for capturing the calibration haemodynamic metrics but can be expanded to provide smooth transitions between phases of the cardiac cycle and additional mechanistic details on arterial dynamics^107^ to better capture the time-course of pressure boundary conditions and to preserve stroke volume balance between the two ventricles.

On the epicardium surface, we model the effect of the pericardium by prescribing a spring Robin type constraint aligned with the surface normals. This constraint, while being able to produce realistic atrioventricular plane displacement, can be extended to more accurately represent frictionless contact mechanics^108^ to better replicate four chamber mechanics^31,49^ and can be important in investigations of pericardial dysfunction^109^.

Finally, our current calibration framework primarily targets left ventricular pressure-volume biomarkers, including volumes, ejection fraction, pressure, and the ECG. Consequently, the model is not explicitly constrained to achieve equal stroke volumes between the right and left ventricles in steady state. This is an important physiological limitation which can be overcome by methods for explicitly tuning right ventricular active tension against measured right ventricle-specific data, as demonstrated by Miller et al.^110^, or a more sophisticated multistep calibration procedure that sequentially optimises circulatory dynamics, passive mechanics and active contraction for both ventricles, as demonstrated by Brown et al.^111^. Both approaches represent a natural extension of the calibration framework presented here, and incorporating independent right ventricular calibration targeting stroke volume balance is a priority for future work. We note that despite the absence of explicit RV calibration, the RV volumetric measures fell within physiological ranges, though the RV peak pressure exceeded the physiological range, consistent with the stroke volume imbalance described above.

## Conclusions

In this study, we show the importance of applying vigorous VVUQ evaluations of ventricular electromechanics for achieving physiological simulations and systematically identify avenues for future work. We set the basis for a strategy for calibrating and validating high-fidelity baseline electromechanical modelling and simulation frameworks. We compiled a comprehensive list of biomarkers for evaluating healthy electromechanical function, and grouped the dataset into non-overlapping calibration and validation sets. We provide evidence of credibility through calibration to haemodynamic and ECG biomarkers and validation by comparison to strain and deformation biomarkers and sensitivity analysis.

Furthermore, through multi-scale sensitivity analysis spanning both mechanical and ECG biomarkers simultaneously, our analyses highlighted the interplay between cellular, tissue, and haemodynamic parameters on the LVEF and provided multi-scale explanations of its link with ECG biomarkers, which is an analysis not previously performed in a fully coupled electromechanical framework. The sensitivity analysis presented here, performed over biologically informed parameter ranges, provides a foundation for future uncertainty quantification studies using probabilistic inference. Taken together, the study provides a systematic framework for credibility assessment of electromechanical cardiac models, identifies open priorities for further development, and represents a meaningful step towards the cardiac Digital Twin vision.

## Acknowledgements

This work was funded in whole, or in part, by the Wellcome Trust (214290/Z/18/Z). For the purpose of Open Access, the author has applied a CC BY public copyright licence to any Author Accepted Manuscript version arising from this submission.

This work was supported by a Wellcome Trust Fellowship in Basic Biomedical Sciences to B.R. (214290/Z/18/Z), the Personalised In-Silico Cardiology (PIC) project, the CompBioMed 1 and 2 Centre of Excellence in Computational Biomedicine (European Commission Horizon 2020 research and innovation programme, grant agreements No. 675451 and No. 823712), the Oxford BHF Centre of Research Excellence (RE/13/1/30181), PRACE-ICEI funding projects icp005, icp013, icp019.

The authors would like to acknowledge Dr Alberto Zingaro and Dr Dimitrios Lialos from ELEM Biotech, who assisted with Alya benchmarking simulations for verification in the revised version of the manuscript.

## Appendix

### 1. Model verification

Following the Land^93^ benchmarks, we generated an ellipsoid geometry and applied pressure and Dirichlet boundary conditions as described in the benchmark. Our simulated results showed good agreement with the mean deformation reported in the original benchmark paper, verifying that the passive mechanical simulations using Alya were free from volumetric locking issues at mesh resolution of ## mm.

**Figure A1.**
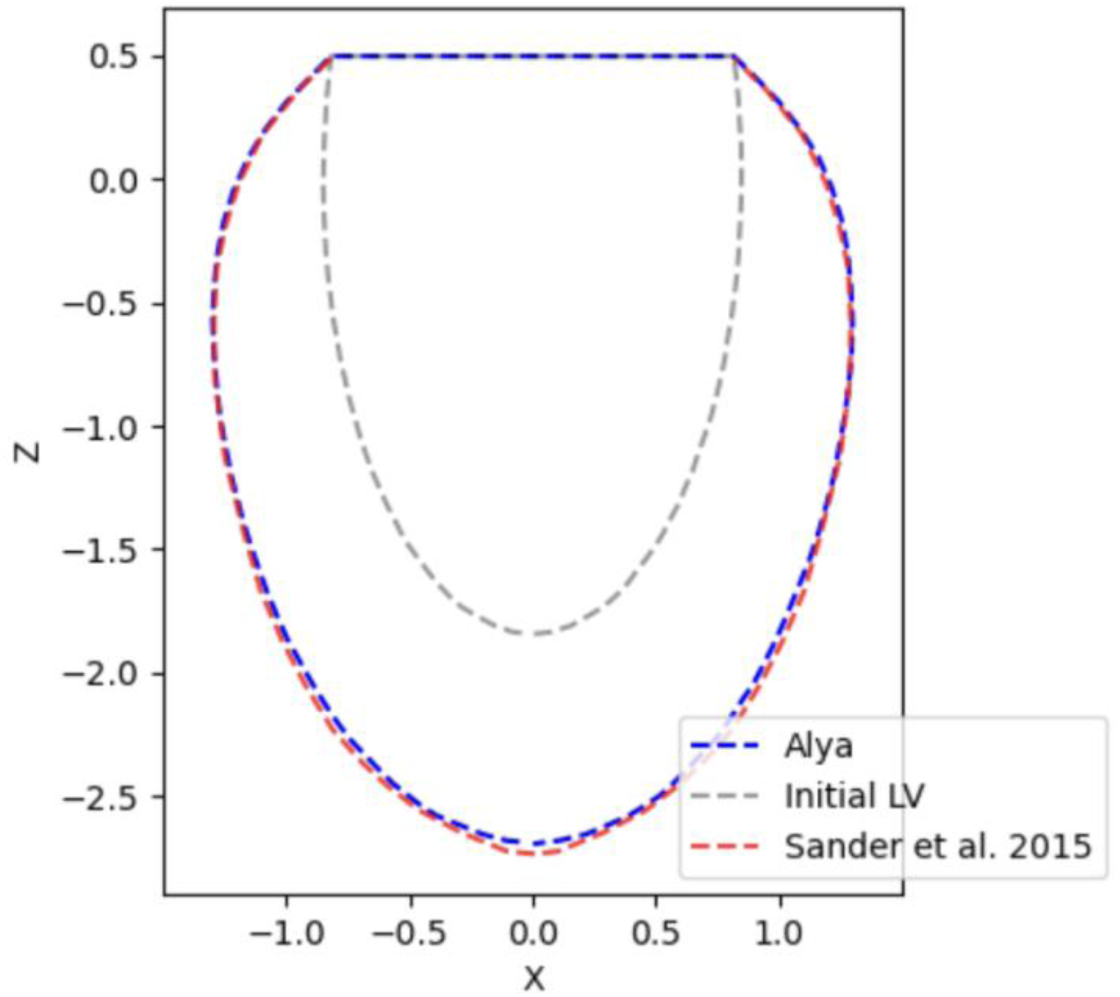
Comparison of mid-line between Alya simulation (grey reference, blue deformed) and Sander Land et al. 2015 (red) passive inflation.

#### Pressure volume calibration strategy design

The goal of the calibration of the initialised set of parameters was to increase LVEF, increase systolic pressure, increase peak filling rate, but decrease peak ejection rate and lower dP/dt_max_. The LVEF and peak systolic pressure had the highest degree of importance for calibration, since they were implicated in a variety of diseases and was well established. The filling rates and dPdt_max_ took a secondary role, since the data came from much smaller sample sizes, and the filling rates were measured using echocardiography techniques, which were not as reliable as volume measurements in CMR (Table 1).

From the sensitivity analysis, we saw that a handful of parameters can increase both LVEF and peak systolic pressure simultaneously: Tref, k_ws_, GCaL, SERCA, Cal50, Kct (Figure 3). However, Tref and kws effects saturate at high values, large changes in SERCA and GCaL can become arrhythmic substrates, Cal50 has a non-monotonic effect on LVEF due to its effect on the diastolic function, and dramatic reductions in Kct brings unrealistic amounts of systolic volume change. A combination of changes in these parameters was needed to dramatically improve LVEF and peak systolic pressure, and in practice, was insufficient by themselves. Decreasing arterial resistance was also needed to help increase LVEF. However, this comes at the cost of reducing peak systolic pressure, which needed to be counterbalanced by increasing the ejection pressure threshold. An increase in ejection pressure increased time spent in isovolumic contraction and isometric force development, and thereby increased the peak systolic pressure. An additional challenge was that there was not a single parameter in our analysis that could decrease peak ejection flow rate (Tref, kws, Kct) and dPdt_max_ (kws) without also decreasing LVEF at the same time. This meant that we needed to achieve higher than healthy LVEF first. With all this in mind, the calibration strategy was as detailed in the Methods section.

### 2. Cellular global sensitivity analysis

**Figure A1:**
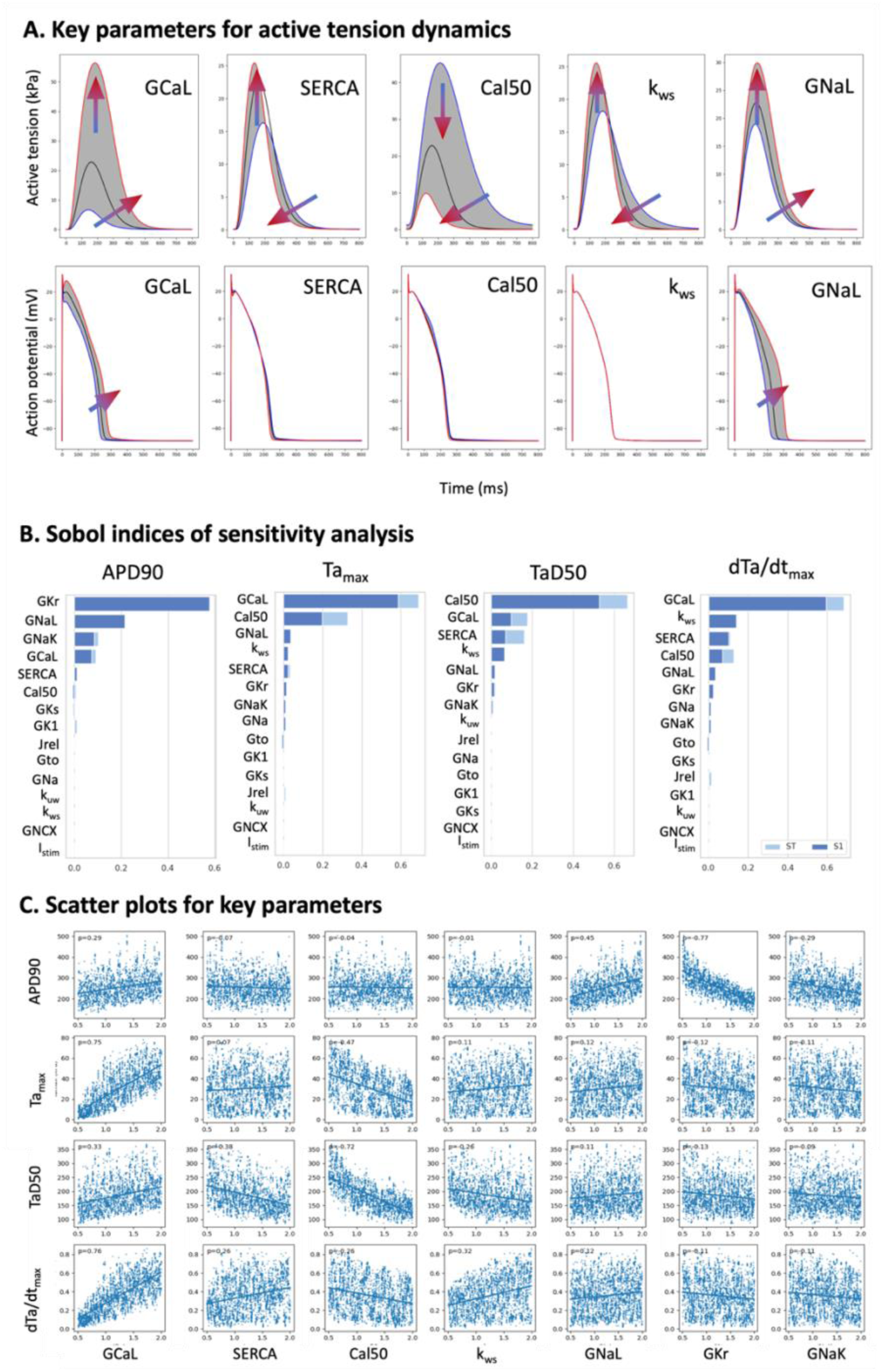
Global sensitivity analysis (C) using cellular model of ventricular electromechanics show the effect of the top five model parameters on active tension (A) and action potential duration (A and B) biomarkers.

### 3. Additional ventricular sensitivity analysis results

**Figure A2.**
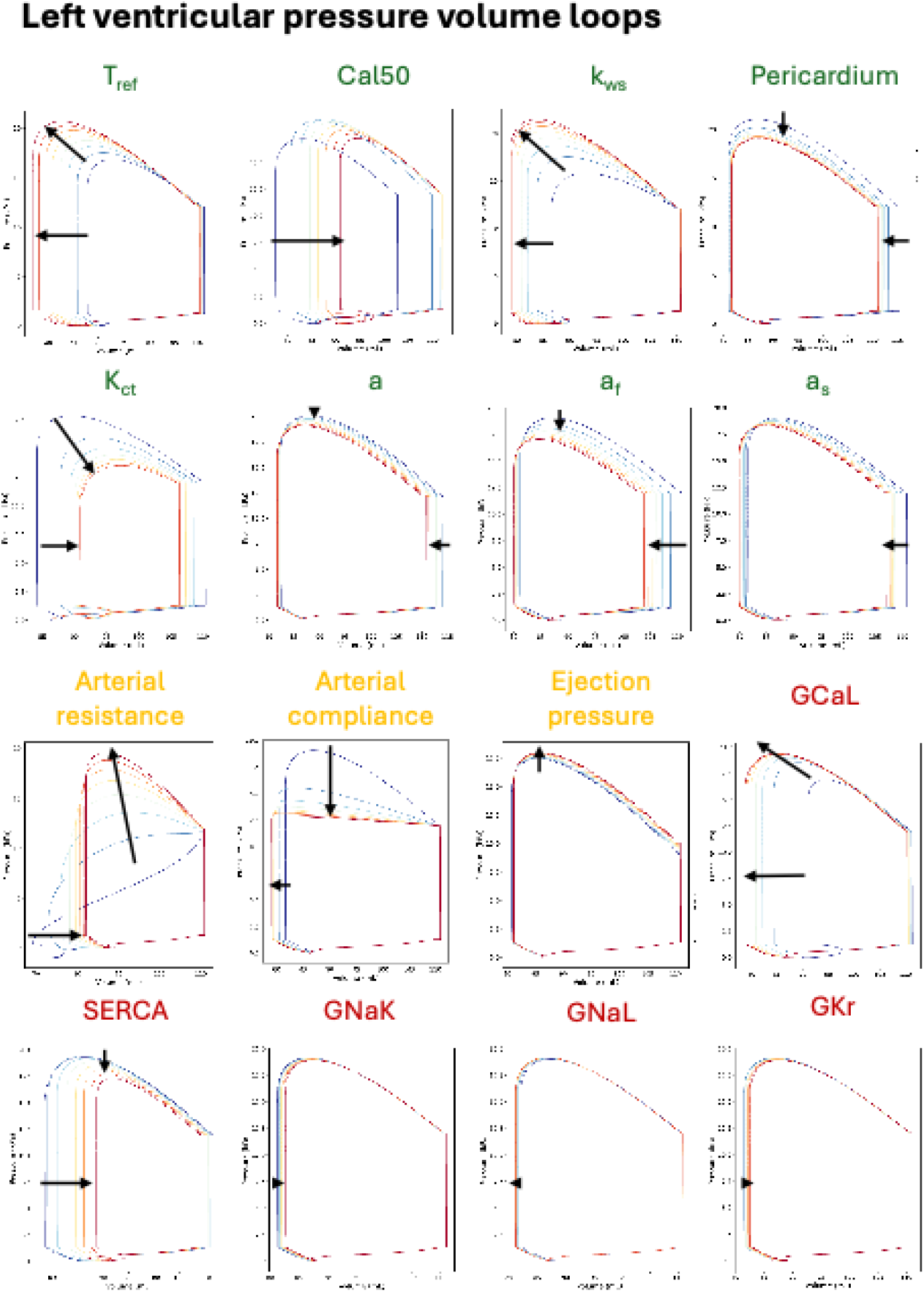
Uncertainties in simulated pressure volume dynamics were influenced by uncertainties in mechanical (green), circulatory (yellow) and ionic conductance (red) parameters of the model.

**Figure A3:**
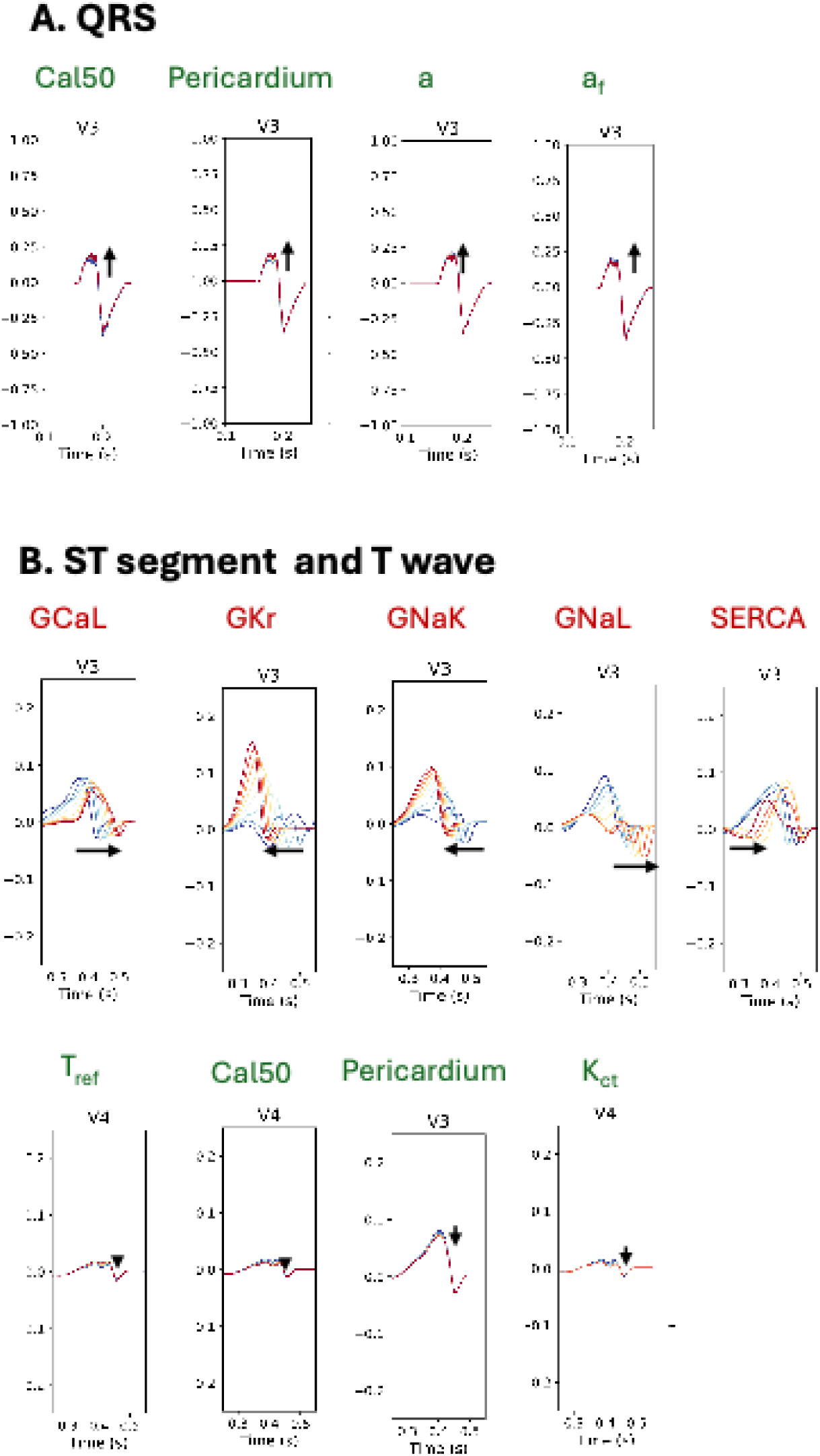
Of the parameters included in the analysis, the QRS section of the ECG showed minor sensitivity to variability in mechanical parameters (green labels) (A) while the ST and T wave segments of the ECG were strongly sensitivity to uncertainties in ionic conductances (red labels), and only showed minor sensitivity to some mechanical parameters (green labels) (B).

**Figure A4:**
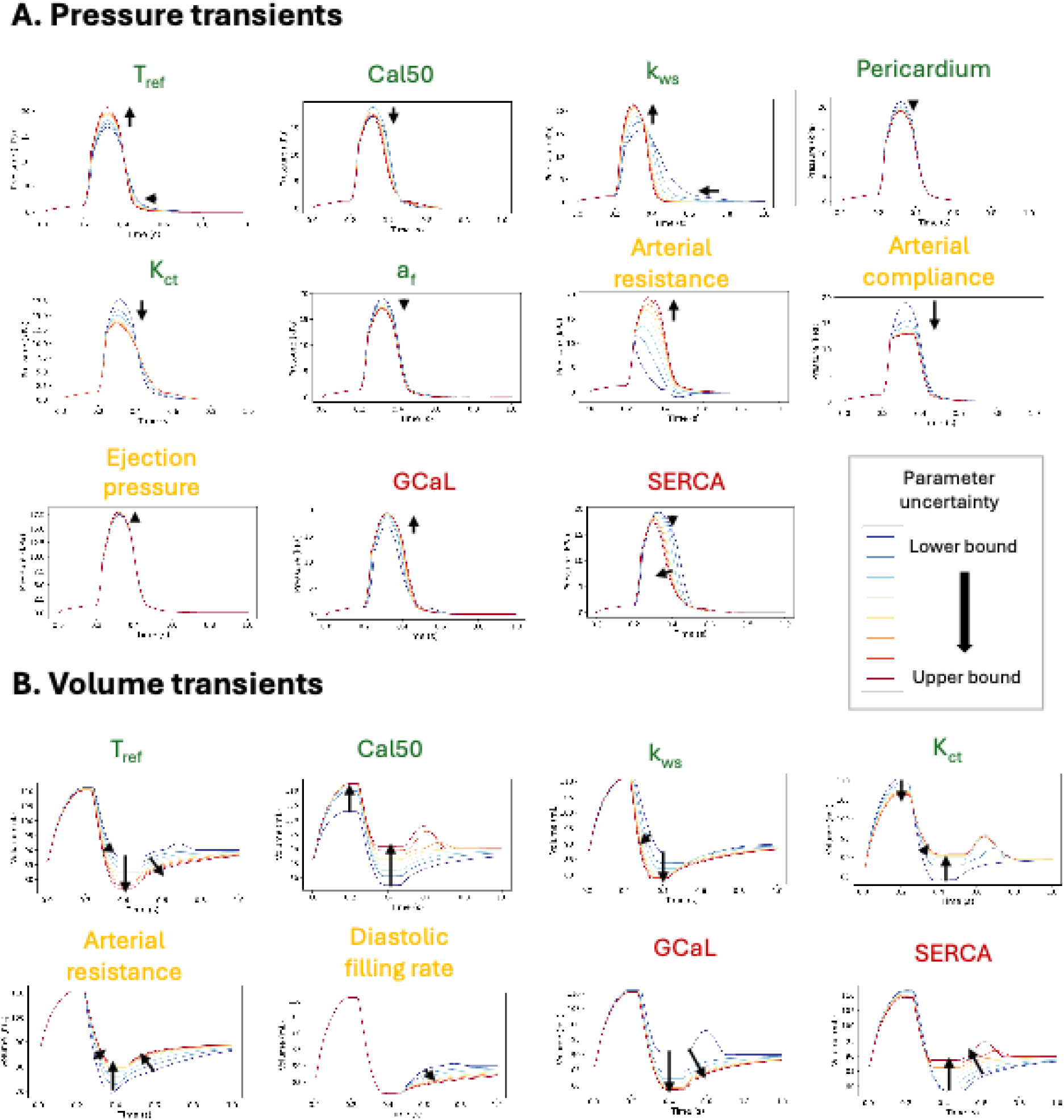
Pressure (A) and volume (B) transients in response to parameter variability in mechanical properties(green), circulation (yellow), ionic conductances (red).

**Figure A5:**
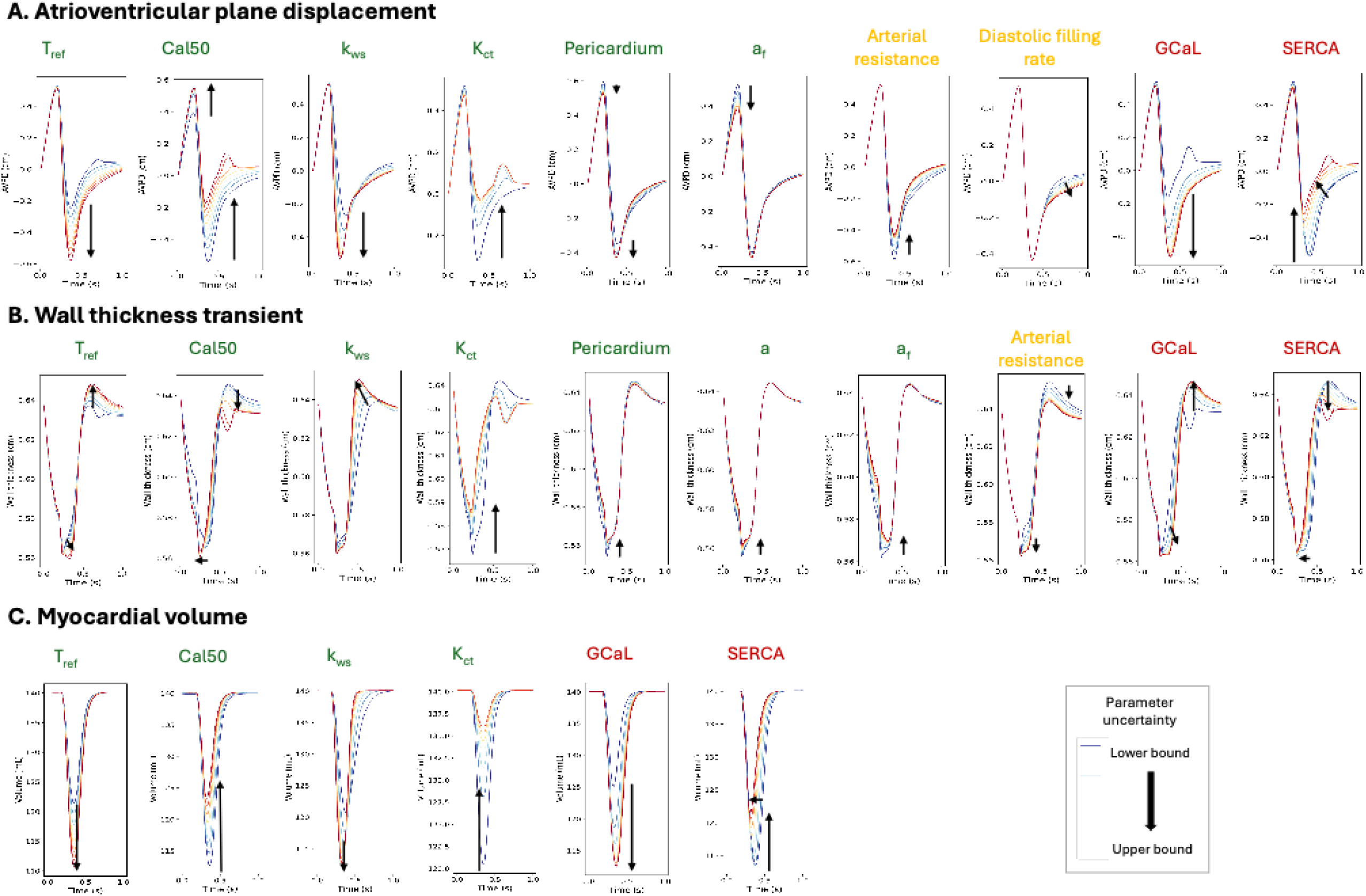
Key aspects of the ventricular deformation, including atrioventricular plane displacement, wall thickness changes, and myocardium volume changes were affected predominantly by mechanical parameters (green) and ionic conductance parameters (red), with weaker effects from circulatory parameters (yellow).

**Figure A6:**
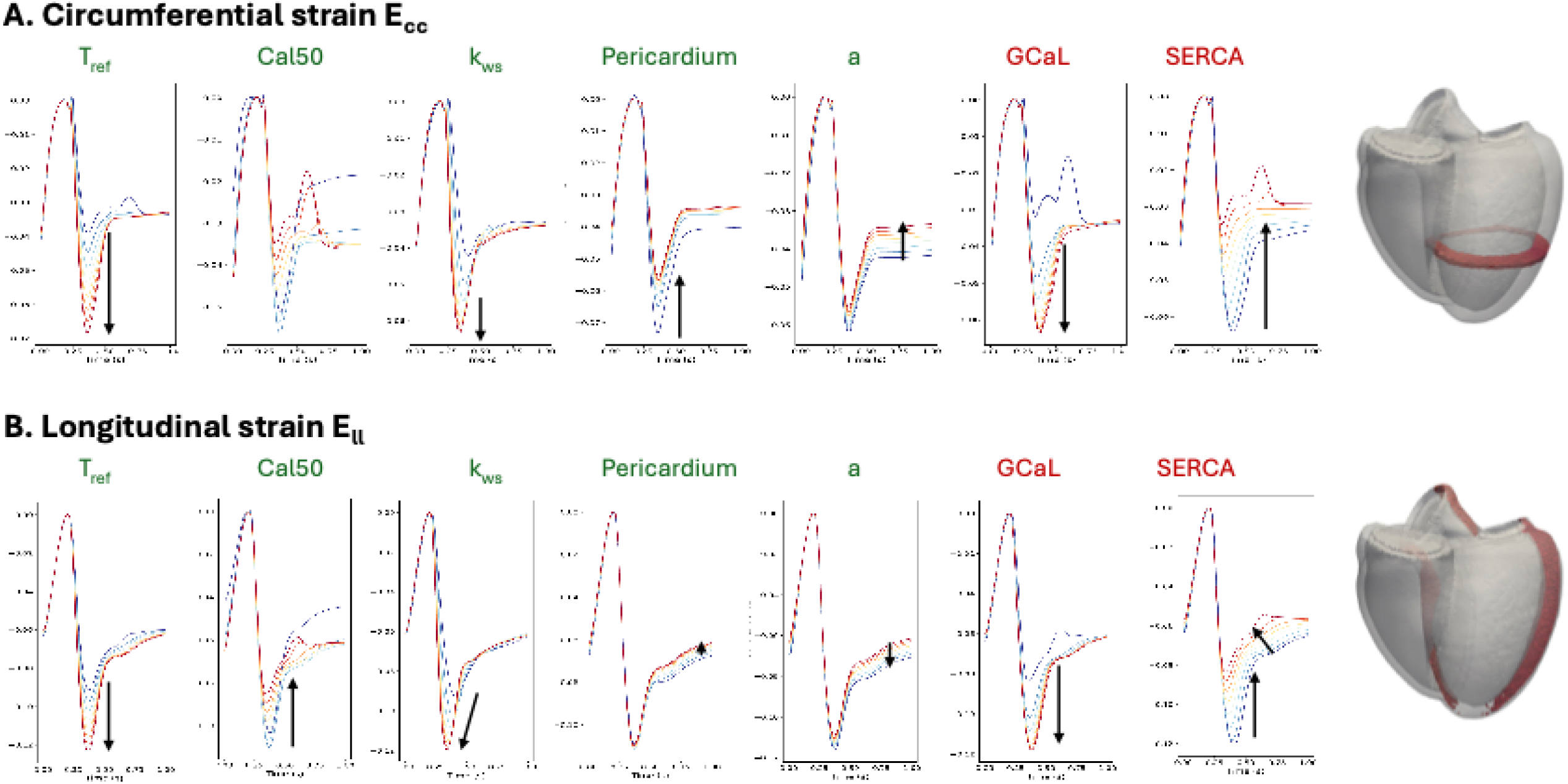
Sensitivity analysis in simulated strain transients were influenced by variability in mechanical (green), circulatory (yellow), and ionic conductance (red) parameters.

